# Carbohydrate active enzymes in *Pectobacteriaceae*: coevolving enzyme sets and host adaptation

**DOI:** 10.64898/2026.05.08.723719

**Authors:** Emma E. M. Hobbs, Tracey M. Gloster, Leighton Pritchard

## Abstract

Many phytopathogenic bacteria have evolved large, diverse arsenals of Carbohydrate Active enZymes (CAZymes) that liberate simple sugars, and thus nutrition and energy, from the complex lignocellulosic matrices of their plant hosts. The CAZyme arsenals of these phytopathogens are expected to be influenced by and adapted to the cell wall composition of their plant hosts. The solutions these organisms have reached for the problem of degrading plant material may help us understand their host ranges and present a rich source of novel CAZymes for exploitation in industrial bioprocessing. Here we catalogue and analyse CAZyme complements (CAZomes) of publicly-available Enterobacterial phytopathogen genomes, including those of the economically significant and widely-studied *Pectobacterium* and *Dickeya* genera. These comprise a broad diversity of CAZymes, providing insight into host adaptation and a resource for bioprospection of industrially-relevant enzymes. We find evidence supporting coevolution of sets of CAZymes specific to bacterial genus and species and, notably, CAZymes associated with pathogen preference for either woody or soft plant tissue, suggesting adaptation of CAZomes to host plant cell wall composition.

## Introduction

Lignocellulose, a complex matrix of structural proteins, aromatic polymers (including lignin) and (ligno)polysaccharides, is the primary component of plant cell walls and one of the most globally abundant renewable carbon resources (1, 2). Industrial bioprocessing of polysaccharide organic materials containing lignocellulose into products such as biofuels, animal feed, and textiles is critical to the development of a greener society and circular economy (3–6). Plant polysaccharides comprise up to 75% lignocellulose by dry weight, primarily as cellulose, hemicellulose and pectin that can be broken down by Carbohydrate Active enZymes (CAZymes) into oligosaccharides and monosaccharides suitable for industrial conversion into bio-based products (1, 7–10). The architecture and chemical composition of lignocellulosic polysaccharides varies between plant tissues and species. Therefore, a large and complex array of CAZymes with diverse substrate specificities is required for efficient, comprehensive industrial degradation, and customised CAZyme formulations are required for some biomass compositions (1, 2, 9, 11–13).

The variable composition of lignocellulose in nature makes design of efficient standard CAZyme mixes for industrial degradation difficult (13). Mixes are often designed by trial-and-error combination of individual enzymes or multiple crude CAZyme mixtures. Alternatively, an organism’s proteome might be screened to identify key CAZyme activities as the foundation of a new mix, subsequently augmented by trial-and-error addition of CAZymes (12, 14–16). Trial-and-error approaches can be slow and resource-intensive, and may optimise the CAZyme mix for a specific feedstock, rather than for broad effectiveness.

Commercially available CAZyme mix inefficiency results in high production costs, low percentage yields, and poor energy efficiency (12, 13, 17, 18). Rationally-designed formulations of targeted CAZyme mixes are proposed as a strategy for improved efficiency but rational design of CAZyme mixes is not widespread (6, 19). Only a minority of CAZymes have been well-characterised and their diversity is not fully understood. Bioinformatic screening of CAZomes from taxonomically and phenotypically diverse plant-degrading microorganisms has been proposed as a means to mine natural variation to enable efficient design of CAZyme mixes, minimising rediscovery of known CAZymes (20).

Many phytopathogenic microbes possess large arsenals of CAZymes that liberate monosaccharides from host plant cell walls in order to fuel growth and/or to penetrate host cells (21, 22). The diversity of plant host lignocellulose architecture is mirrored by that of of phytopathogen CAZomes, positioning phytopathogens as a potentially rich source of industrially exploitable CAZymes (9, 23).

Suppression and evasion of host defence responses is essential for pathogen survival and is known to place strong selective pressures on a pathogen’s effector complement. The same evolutionary “arms race” between pathogen and host may also drive pathogen CAZome diversity (24). Degradation of host cells by a pathogen may trigger host damage-sensitive response mechanisms that inhibit pathogen progress, so CAZymes may be under selection pressure to avoid triggering Damage-Associated Molecular Pattern (DAMP) recognition, resulting in variation of CAZome composition by host (19, 20, 25–27).

Roles for CAZymes in disease interactions are known *via* single species or single isolate interactions with a host, but there have been few systematic comparisons of CAZome variation at genus or taxonomic Family level (20, 28–30). Widening comparisons to Family level may reveal new associations between CAZome composition, host range, environmental niche, phenotype, and lineage. An ability to relate CAZome complement to host range could potentially enable prediction of host-jump potential and improve modelling of impacts of emerging plant pathogens, with benefits to food security and regulation.

If CAZymes act concertedly as evolutionarily optimised groups of complementary enzymes to break down complex plant material, it should be feasible to detect co-evolution of CAZyme sets targeting specific substrates. Such co-evolving CAZyme sets could be useful leads in rational design of optimal enzyme mixes for industrial degradation of lignocellulosic biomass (20, 28).

CAZy (www.cazy.org) is the most authoritative and comprehensive CAZyme bioinformatic resource. The CAZy database formally classifies CAZymes into six broad functional classes by biochemical activity: (i) glycoside hydrolases (GHs); (ii) glycosyltransferases (GTs); (iii) polysaccharide lyases (PLs); (iv) carbohydrate esterases (CEs); (v) auxiliary activities (AAs); and (vi) non-catalytic carbohydrate-binding modules (CBMs). Each CAZy class is further sub-divided into sequence-based families corresponding to presumed shared mechanisms and structural folds. Class and family-level classifications enable principled annotation and comparison of CAZome biochemical and functional diversity (29, 31).

Aerobic fungi (especially *Trichoderma* and *Aspergillus*) have been exploited to formulate commercial lignocellulolytic CAZyme mixes (13, 19). By contrast, other plant pathogen-rich lineages are relatively underexplored in this context, such as the *Pectobacteriaceae* Family (NCBI:txid1903410). This family encompasses Gramnegative, rod-shaped bacteria distributed across the genera *Acerihabitans* (NCBI:txid2811373) (32), *Affinibrenneria* (NCBI:txid2853326) (33), *Brenneria* (NCBI:txid71655) (34), *Dickeya* (NCBI:txid204037) (35), *Lonsdalea* (NCBI:txid1082702), *Musicola* (NCBI:txid2884243) (36), *Pectobacterium* (NCBI:txid122277) (37), and *Samsonia* (NCBI:txid160433) (36, 38). The *Pectobacteriaceae* include several economically important food crop pathogens, including *Pectobacterium carotovorum* and *Dickeya dianthicola*, which can reduce yields by up to 45% (39–42). *Pectobacteriaceae* phytopathogens are distributed globally and cause disease on a diverse range of plant hosts, across both monocots and dicots (43). However, *Pectobacteriaceae* genera can also be divided broadly by host tissue preference into hard or woody plant tissue (e.g. bark and nuts) degraders, such as *Acerihabitans, Affinibrenneria, Brenneria, Lonsdalea, Samsonia*, and *Sodalis*, or soft plant tissue (e.g. stem and leaves) degraders - also known as soft rot *Pectobacteriaceae* (SRP) - including *Pectobacterium* and *Dickeya* (32, 44–46).

Significant genomic diversity across the *Pectobacteriaceae* family has provided a basis for recent taxonomic reclassifications (36, 38). However, to our knowledge, no genomic explanation for the distinct soft vs hard plant tissue targeting phenotypes within this Family has yet been identified.

The most important virulence characteristic shared by *Pectobacteriaceae* is considered to be secretion of large plant cell wall degrading CAZyme arsenals (46, 47). We therefore hypothesise that not only may *Pectobacteriaceae* CAZomes be a rich source of potentially industrially exploitable CAZymes, but differences between their composition may also be associated with the distinct host ranges of *Pectobacteriaceae* species, and specifically their ability to exploit hard or soft host plant material.

To assist in our analysis, we developed `cazomevolve`(‘cazome-evolve’, DOI:10.5281/zenodo.6614827), a software package for taxonomically-aware exploration and comparison of CAZomes that identifies associations between metadata classes (such as niche or phenotype) and CAZome composition. We demonstrate the application of `cazomevolve`by exploring the prevalence of CAZy families in *Pectobacteriaceae* genomes, and their association with host range. We (i) compare predicted CAZome compositions of 717 *Pectobacteriaceae* genome sequences obtained from GenBank; (ii) identify CAZy families that co-occur with potential to form the foundation of a minimal enzyme mix for lignocellulose degradation; (iii) identify associations between CAZy families and host range (and, more generally, soft or hard plant tissue preference); and (iv) for the first time (to our knowledge), use `coinfinder`(48) to identify lineage-specific networks of co-evolving CAZy families that could inform rational design of industrial enzyme mixes.

## Methods

We used the software package `cazomevolve` (version v0.1.5) (DOI:10.5281/zenodo.6614827) to carry out analyses in this manuscript. Detailed description of `cazomevolve`, including a summary of its implementation, is presented in Supplementary Information.

All bash, R, and Python scripts and commands used in this manuscript are provided in the GitHub repository https://hobnobmancer.github.io/SI_Hobbs_et_al_2024_Pecto/ (DOI:10.5281/zenodo.7699656), including a README file walkthrough, high resolution figures, and additional analyses not presented in this manuscript. Manuscript figures were generated using the Jupyter note-book `explore_pectobact_cazomes.ipynb`, also available in the GitHub repository.

Default settings were used for all software tools unless specified. Complete parameter sets are provided in accompanying scripts. All analyses for the manuscript were performed on a PC with an Intel Xeon E3-1200 v2 (Ivy Bridge, IBRS) processor, 64GB RAM, running Ubuntu 16.04.5 LTS.

All supplementary figures and tables are compiled into a PDF file, which is available at the GitHub repository: https://github.com/HobnobMancer/SI_Hobbs_et_al_2024_Pecto/blob/master/Hobbs_et_al_SI_Pectobacteriaceae.pdf. A visual summary of these methods is presented in SI figures 1-4.

### Downloading and annotating *Pectobacteri-aceae* genomic assemblies

The bash script `download_ms_genomes.sh`was used to configure `cazomevolve`to query the NCBI Assembly database (49) with the query term “Pectobacteriaceae”, downloading genome (.fna) and proteome (.faa) FASTA files for 717 GenBank assemblies associated with NCBI:txid1903410 (queried 23rd April 2023).

The Python script `ident_missing_proteomes.py`was used to identify assemblies where a proteome FASTA (.faa) file was not retrieved from NCBI. In these cases, the genome was annotated using Prodigal (v2.6.3) (50), co-ordinated by the `annotate_pectobact_genomes.sh`bash script. The resulting combined set of protein sequence annotations (NCBI and Prodigal) is referred to as the *Pectobacteriaceae* dataset.

A visual overview of how the proteome dataset was compiled for each input genome is presented in SI figure 1.

### Construction of a local CAZyme database

All CAZyme records were downloaded from CAZy on 20th January 2023 and compiled into a local database using `cazy_webscraper`(version 2.2.3) (51), configured using the script `build_cazyme_database.sh`.

### CAZome annotation

The bash script `get_cazy_cazymes.sh`was used to configure `cazomevolve`to import existing CAZy annotations (discarding subfamily numbers) for each genome from the local CAZy database into the expanded CAZyme database. Proteins from each genome that had no annotation in CAZy were submitted to the CAZyme classifier dbCAN v2.0.11 (52) using `cazomevolve`. The analysis was configured by bash scripts `run_dbcan.sh`and `get_dbcan_cazymes.sh`to compile a comprehensive CAZyme complement (i.e. CAZome) for each genome, combining dbCAN and CAZy annotations for each genome into a single reference CAZyme database (the “expanded CAZyme database”) using `cazomevolve`.

A visual overview of this approach is presented in SI figure 2.

### NCBI taxonomic classification

The bash script `add_taxs.sh`, was used to download taxonomic assignments from the NCBI Taxonomy database for each genome accession in the expanded CAZyme database (SI figure 3).

### Average nucleotide identity (ANI)

The `pyani`v0.3.0- alpha software, configured by the script `run_pyani.sh`, was used to calculate pairwise average nucleotide identity (ANIm) for *Pectobacteriaceae* genomes (53) (SI figure 4). The R script `build_ani_tree.sh`was used to generate a Newick format dendrogram from the ANI identity matrix, rendered using the R script `build_anim_tree.R`using the R packages ape v.5.7.1 (54) and phytools v.2.0.3 (55). The Python script `add_ani_tax.py`was used to annotate the tree with taxonomic data.

### Exploration and visualisation of CA-Zome composition

The Jupyter notebook `explore_pectobact_cazomes.ipynb`was used to compare CAZome size and composition in the *Pectobacteriaceae* genomes, using `cazomevolve.explore`, as summarised below. All statistical tests were conducted in R using StatsModels v0.14.4(56) with a significance threshold of P<0.05 unless otherwise specified.

#### Comparison of proteome and CAZome sizes

Summary statistics were generated across all genomes and per genus, recording (i) CAZymes per genome; (ii) total proteins per genome; (iii) CAZome proportion of the proteome; and (iv) count of CAZyme families. For each variable, one-way ANOVA was used to test whether means varied (i) across all genera, (ii) across soft plant tissue-degrading (PTD) genera (*Pectobacterium* and *Dickeya*), hard PTD genera (*Acerihabitans, Affinibrenneria, Brenneria, Lonsdalea and Sodalis*), and *Musicola* (this genus has unclear preference for soft or hard plant tissue (36)). Where a statistically significant difference was found, a post hoc Tukey HSD test was used to identify contrasts where means differed significantly (57).

#### CAZyme class frequencies

The frequency of each CAZyme class was calculated as the count of unique protein accessions annotated with at least one CAZyme family in that class. Mean and standard deviation of each class frequency was calculated for each CAZyme class and genus. A proportional area plot of mean CAZyme class frequency per genus was made using RAWGraphs v2.0 (58). We used two-way ANOVA to test whether mean CAZyme class frequency varied across CAZyme class and genus as main effects, and also for evidence of CAZyme class-genus interaction. Within each individual CAZyme class, one-way ANOVA was used to determine if mean frequencies varied across genera, followed by Tukey HSD test to identify genus contrasts with significant pairwise differences. One-way ANOVA and post hoc Tukey HSD tests were also used to test for evidence of a difference in means between soft and hard plant tissue targeting genus groups.

#### CAZyme family frequencies

A count of unique protein accessions associated with each CAZyme family was calculated for each genome (the CAZyme family frequency). A clustermap of CAZyme family frequencies was generated using the Seaborn package (v0.12.2) (59). CAZyme families present in all genomes of a group (e.g. a single genus) were collectively considered to be the ‘core CAZome’ for that group. Core CAZomes were determined for the full dataset, and for each genus. CAZyme families present in only a single genus (i.e. genus-specific families) were also identified.

#### Principal component analysis

Principal component analysis (PCA), implemented in scikit-learn v1.2.1 (60), was applied to a table of CAZyme family frequencies for each genome using `cazomevolve.explore`.

#### Rarefaction curves and CAZome completeness estimation

To estimate the expected total number of unique CAZyme families in each genus and the completeness of current sampling, we constructed rarefaction curves by iterative resampling of the genomes in each genus, recording the number of new CAZyme families discovered with each sampled genome. 100 replicate runs were performed per genus and the resulting average curves, with variance indicated as a ribbon showing the complete range of individual curves, were plotted in Python using Seaborn v0.12.2 (59).

#### CAZyme family co-occurrence

We used `cazomevolve.explore`to identify groups of co-occurring families that were always found to be annotated as present together in a defined group of genomes. Co-occurring families were found for the complete dataset, separately for each genus, and for the hard- and soft-tissue-targeting groups. The number of genomes in which co-occurring families were present together was computed as the *incidence* of that collection of families, and plotted with UpSetPlot v0.9.0 (61).

### Construction of dendrograms and tanglegrams to compare distance-based trees

Dendrograms were built from CAZyme family frequencies and pyani ANIm datasets using Euclidean distance and single hierarchical clustering with the R script `build_tanglegrams.R`. The dendrograms were ladderised, and each leaf branch colour coded by genus, then they were used to construct a tanglegram using dendextend v1.18 (62).

### Identification of co-evolving CAZyme family networks

coinfinder v1.0.9 (48) was used to identify groups of CAZyme families that appear in the same genome together more often than expected by chance, taking into account lineage effects implied by the ANI dendrogram. The software was configured using the bash script `find_coevolving_pectobact.sh`and the bash script `add_taxs_tree.sh`was used to annotate the dendrogram with NCBI taxonomic classifications .

## Results

All datasets and full-sized figures are available in the GitHub repository https://hobnobmancer.github.io/SI_Hobbs_et_al_2024_Pecto/. Supplementary (SI) figures and tables are available in the GitHub repository at https://hobnobmancer.github.io/SI_Hobbs_et_al_2024_Pecto/Hobbs_et_al_SI.pdf.

### CAZymes comprise approximately 2.5% of the *Pectobacteriaceae* proteome

Genome and proteome sequence data were downloaded for 717 *Pectobacteriaceae* publicly-available GenBank genome assemblies at NCBI. No proteome data was available for 107 assemblies, so these genomes were annotated using `Prodigal`. A local CAZyme database was constructed using `cazy_webscraper`, containing all CAZyme records from the CAZy database (http://www.cazy.org) (51). All 2,994,018 *Pectobacteriaceae* proteome protein IDs were queried against this database using `cazomevolve`, identifying CAZyme family annotations for 17,132 (0.06%) records from 160 (22%) of the 717 publicly available genomes. The remaining proteome sequences were provided as input to `dbCAN`, which predicted an additional 60,994 CAZyme sequences, for a total of 78,132 CAZymes (2.6% of the predicted *Pectobacteriaceae* proteome). Table 1 summarises proteome size, counts of CAZyme families, and CAZymes per genome in *Pectobacteriaceae*, and visual summaries of the data are presented in SI figures SI5-SI8.

**Table 1.**
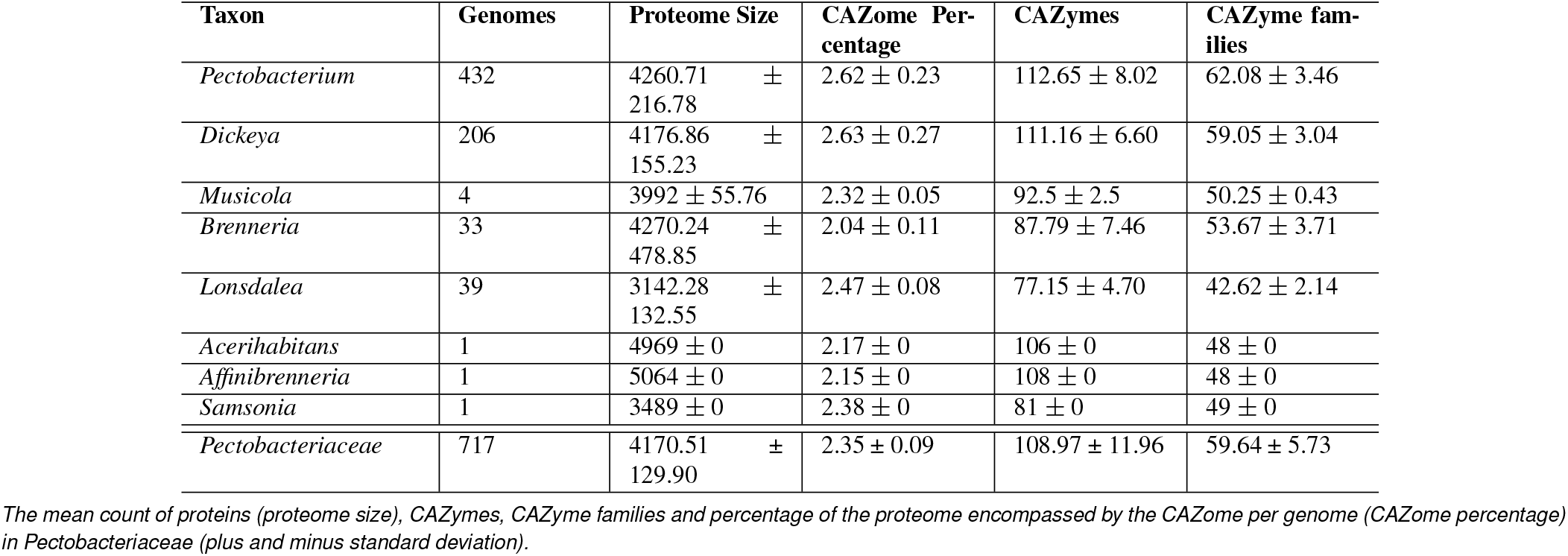
Summary of *Pectobacteriaceae* CAZome sizes.

### Absolute and relative CAZome sizes are not uniform across *Pectobacteriaceae*

#### Proteome sizes vary between genera, but do not clearly distinguish between soft/hard tissue degradation

One-way ANOVA indicates that mean proteome size varies across *Pectobacteriaceae* genera (one-way ANOVA, P=1.6E-130) (SI figure 5). Proteome sizes of *Acerihabitans* and *Affinibrenneria* are significantly larger than those of other genera (SI table 1, SI figure 5) but there is only one representative genome for each of these, limiting generalisation. *Lonsdalea* and *Samsonia* have smaller proteomes than other genera (though *Samsonia* and *Musicola* have similar proteome sizes). *Pectobacterium, Dickeya, Brenneria* and *Musicola* have similar mean proteome sizes, but data do not support a simple association between proteome size and soft or hard plant tissue degrading phenotype.

#### CAZymes are a larger proportion of the proteome in SRP than HTD pathogens

The proportion of the proteome made up of CAZymes varies by host range category (one-way ANOVA, P=7.9E-77, SI figure 6). The SRP proteome comprises a larger proportion of CAZymes compared to *Musicola* (Tukey HDS, P=0, mean difference=-0.3) and HTD genomes (Tukey HSD, P=0, mean difference=-0.4) (SI table 2). The CAZome proportion of the proteome also varies across *Pectobacteriaceae* genera (one-way ANOVA, P=1.6E-101).

#### SRP CAZomes are larger than HTD CAZomes

CAZome size varies across *Pectobacteriaceae* genera (one-way ANOVA, P=3.3E-142, SI figure 7). *Pectobacterium* and *Dickeya* have the largest CAZomes (excluding *Affinibrenneria* and *Acerihabitans*). Mean CAZome size was found to vary between SRPs, HTDs, and *Musicola* (one-way ANOVA, P=3E-135, Tukey HSD of all contrasts P *≤* 0.04, SI table 5) such that SRP genomes have the largest mean CAZome size (Tukey’s HSD, P=0, mean difference=29).

#### SRP CAZomes contain a larger number of CAZyme families

The mean number of CAZy families differs between *Pectobacteriaceae* genera (one-way ANOVA, p=2.6E-167, SI figure 8). *Pectobacterium* and *Dickeya* CAZomes have significantly more CAZy families than *Musicola* and any of the HTD genera (Tukey’s HSD P *≤* 0.002, except for the contrast between *Dickeya* and *Samsonia*: P=0.05). *Pectobacterium* has more CAZyme families than *Dickeya* (P=0, SI table 6). The mean number of CAZy families per genome varies between SRP, HTD, and *Musicola* genomes (one-way ANOVA, P=2E-116). SRPs have more CAZy families than *Musicola* (Tukey HSD, P=0) and HTD genomes (Tukey HSD, P=0) and there is no difference in mean number of CAZyme families between *Musicola* and HTD genomes (Tukey HSD, P=0.4). (SI table 7).

Taken together these results indicate that CAZome composition differs significantly across the *Pectobacteriaceae* genera. Notably, in SRPs a larger proportion of the proteome is taken up by the CAZome, which also contains on average more CAZymes and more CAZyme families, than in HTDs or *Musicola. Musicola*’s CAZome composition is more similar to HTDs than SRPs. We interpret these results to reflect that CAZome composition may broadly correlate with niche adaptation for SRPs and HTDs, although lineage may also contribute to CAZome size and diversity.

#### CAZy classes are differentially represented in pathogens that degrade soft and hard plant tissue

We calculated (i) the count per genome (frequency) of CAZymes from each CAZy class and (ii) the percentage of the CAZome represented by each CAZy class per genus across all *Pectobacteriaceae* (figure 1). One-way ANOVA showed that mean frequency varies across genera for each CAZy class (table 2). Two-way ANOVA indicated that frequency of CAZy class representation is varies with CAZy class, genus, and the interaction between the two (SI table 8).

**Fig. 1.**
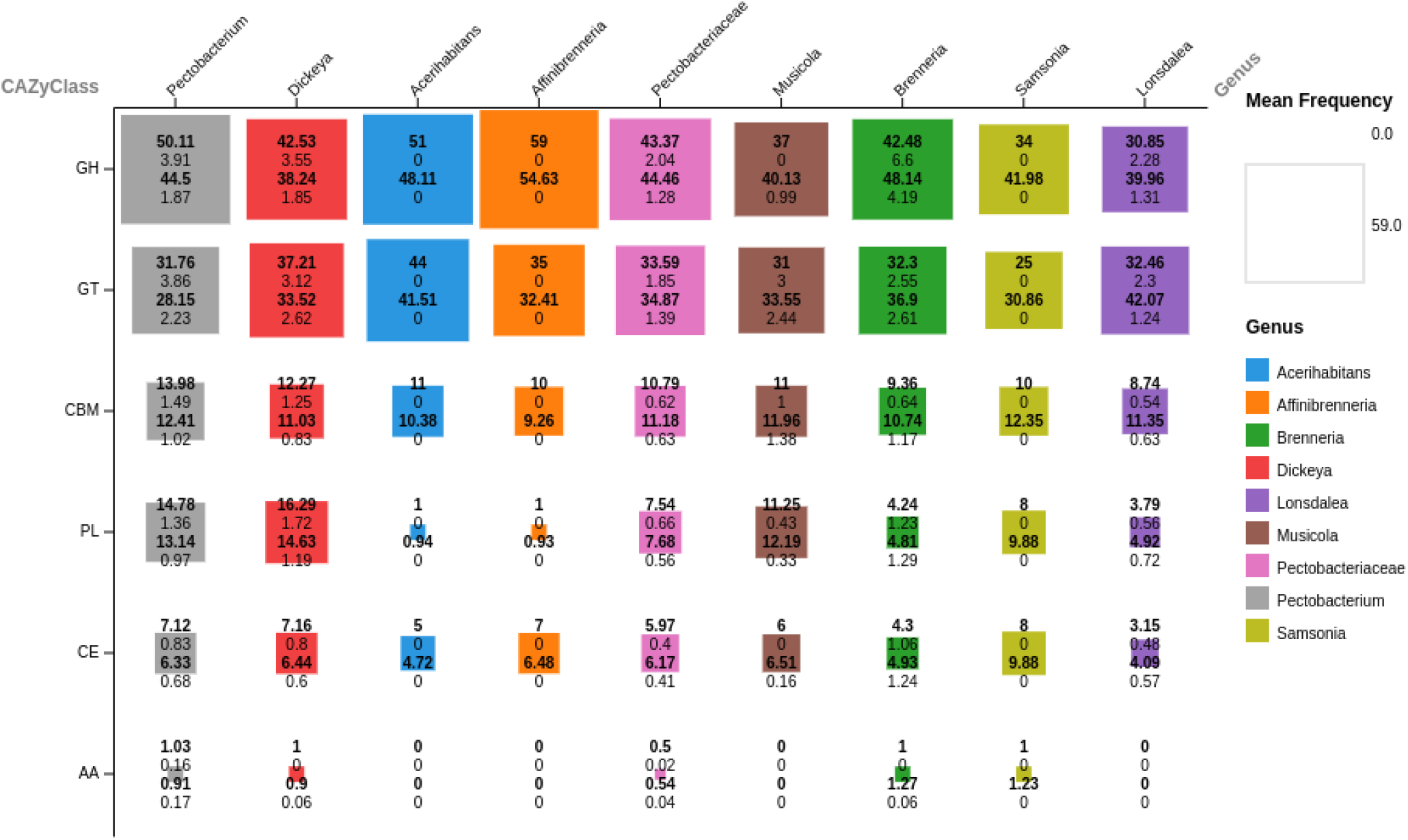
Proportional area plot of the mean frequency of CAZymes per CAZy class in *Pectobacteriaceae*. Each square is annotated (top to bottom) with: mean frequency (bold), frequency standard deviation, mean proportion of the CAZome (%, in bold), standard deviation of proportion.

**Table 2.**
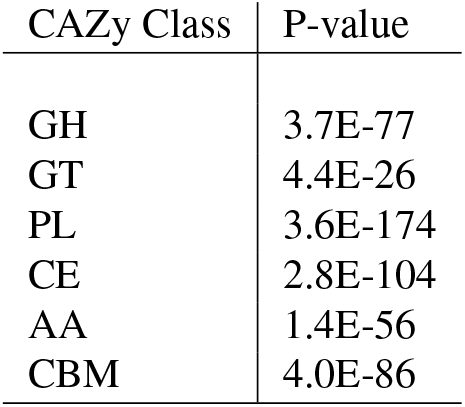
One-way ANOVA results for CAZy class frequency across *Pectobacteriaceae* genera. *p*-values *≤* 0.05 indicate that mean CAZy class frequency is not uniform across all genera.

| CAZy Class | P-value |
| --- | --- |
| GH | 3.7E-77 |
| GT | 4.4E-26 |
| PL | 3.6E-174 |
| CE | 2.8E-104 |
| AA | 1.4E-56 |
| CBM | 4.0E-86 |

The majority of the CAZome for each genus comprises GH (mean 44% ± 1 SD) and GT family members (mean 35% ± 1 SD) (figure 1). *Affinibrenneria* and *Acerhabitans* contained the largest mean count of GH and GT enzymes, respectively. *Lonsdalea* genomes contain significantly fewer GH enzymes than all genera except *Musicola* and *Samsonia*

(Tukey’s HSD, P>0.05). *Pectobacterium* CAZomes contain significantly more GHs than *Brenneria, Dickeya* and *Musicola* (Tukey’s HSD, P *≤* 0.0006, SI tables 9 and 10). *Lonsdalea* CAZomes contain fewer CE enzymes than all other genera except *Acerhabitans* (P *≤* 0.01, SI table 11). *Brenneria* CAZomes contain a larger proportion of CE enzymes than *Dickeya, Pectobacterium* and *Samsonia* (Tukey’s HSD P *≤* 0.02). We find no clear association of GH, GT, and CE CAZyme class distribution with SRP or HTD phenotype.

#### PL, AA, and CBM class representation is expanded in SRP *Pectobacteriaceae*

The mean frequency of CBM domains in each CAZome was found to be broadly comparable across the genera, with highest frequency seen in *Pectobacterium* and *Dickeya* genomes (figure 1). However, we found statistically significant differences between SRP and HTD groups (one-way ANOVA, p-value=1.7E-65), between *Pectobacterium* and *Musicola* (Tukey HSD, P *≤* 0.01; SI table 12), and between SRPs, HTDs, and *Musicola* (Tukey HSD, P*≤*0.02, SI table 13) .

The Auxiliary Activity (AA) class is the least well-represented across *Pectobacteriaceae*, present in only 465 (65%) of the genomes studied. AA CAZymes are found in 41% of *Dickeya* and 24% of *Brenneria*, but in 84% of *Pectobacterium* genomes (83.6%). Most *Pectobacteriaceae* genomes possess a single AA enzyme, but ten *Pectobacterium* genomes contain two. SRP genera contain more AA enzymes than *Musicola* or HTD genera (One-way ANOVA p-value=3.1E-25; Tukey HSD; SI table 15).

Polysaccharide lyase (PL) class members were found at greater frequency in the SRP genera *Pectobacterium* and *Dickeya*, and in *Musicola* (11-16 representatives on average), than in hard PTD genera (1-4 representatives on average, not including *Samsonia*) (figure 1) (Tukey’s HSD, P<0.05; SI table 16).

Only one *Samsonia* genome (*S. erythrinae*) was available, and this contains more PLs than are seen in other HTD genera. However, *Samsonia* shares a more recent ancestor with *Pectobacterium* than with the other HTD genera, and this observation may reflect a lineage effect (63).

Plant pathogen PLs are expected to target pectin and pectate, the lignocellulosic polysaccharides most abundant in soft plant tissues (64, 65). Adaptation to soft plant tissue is consistent with statistically significant overrepresentation of PL members in SRP genera and *Musicola* (Tukey’s HSD, P *≤* 0.003; SI table 16). Taken together, our observations are consistent with phenotype-specific expansion of PL, AA and CBM CAZyme domains in soft tissue-degrading *Pectobacteriaceae*, suggesting an association between niche and CAZome composition.

#### Variation in CAZome composition may be explained by preference for soft and hard plant tissue, and by lineage

We interpret the described variation in CAZome composition and expansion of CAZy classes in specific genera to suggest that: (i) variation in the CAZome may result by adaptation of pathogens to a lifestyle involving soft or hard plant tissue degradation; and (ii) that *Pectobacteriaceae* CAZome composition might allow genomes to be clustered into their respective genera, reflecting the combined influence of lineage and host adaptation. We used Principal Component Analysis (PCA) of each CAZome’s CAZy family representation to investigate whether there was evidence to support this interpretation.

Dimensional reduction by PCA of CAZy family frequencies captures variance in the data set in principal components as described in a scree plot (PCs, SI figure 9a). The first four components - PC1 (15%), PC2 (12%), PC3 (6%), and PC4 (5%) - capture approximately 40% of total variance, but substantially more than any other individual PC (SI figure 9b). We projected the *Pectobacteriaceae* CAZomes onto these four PCs (figure 2) colouring by genus and by species (individual score and loadings plots are available in the online repository). A scores plot of PC1 and PC2 (figure 2) gathers the CAZomes into clusters that correspond well to genus. The ordering of genomes along PC1 forms a ranking from soft to hard plant tissue targeting phenotypes: SRP *Pectobacterium* and *Dickeya* CAZomes cluster at values near zero and on the negative axis of PC1. Conversely, all hard plant tissue targeting genera are clustered in the positive PC1 axis. We interpret this to mean that soft/hard tissue phenotype is an influential factor that explains a substantial component of variation in CAZome composition across *Pectobacteriaceae*.

**Fig. 2.**
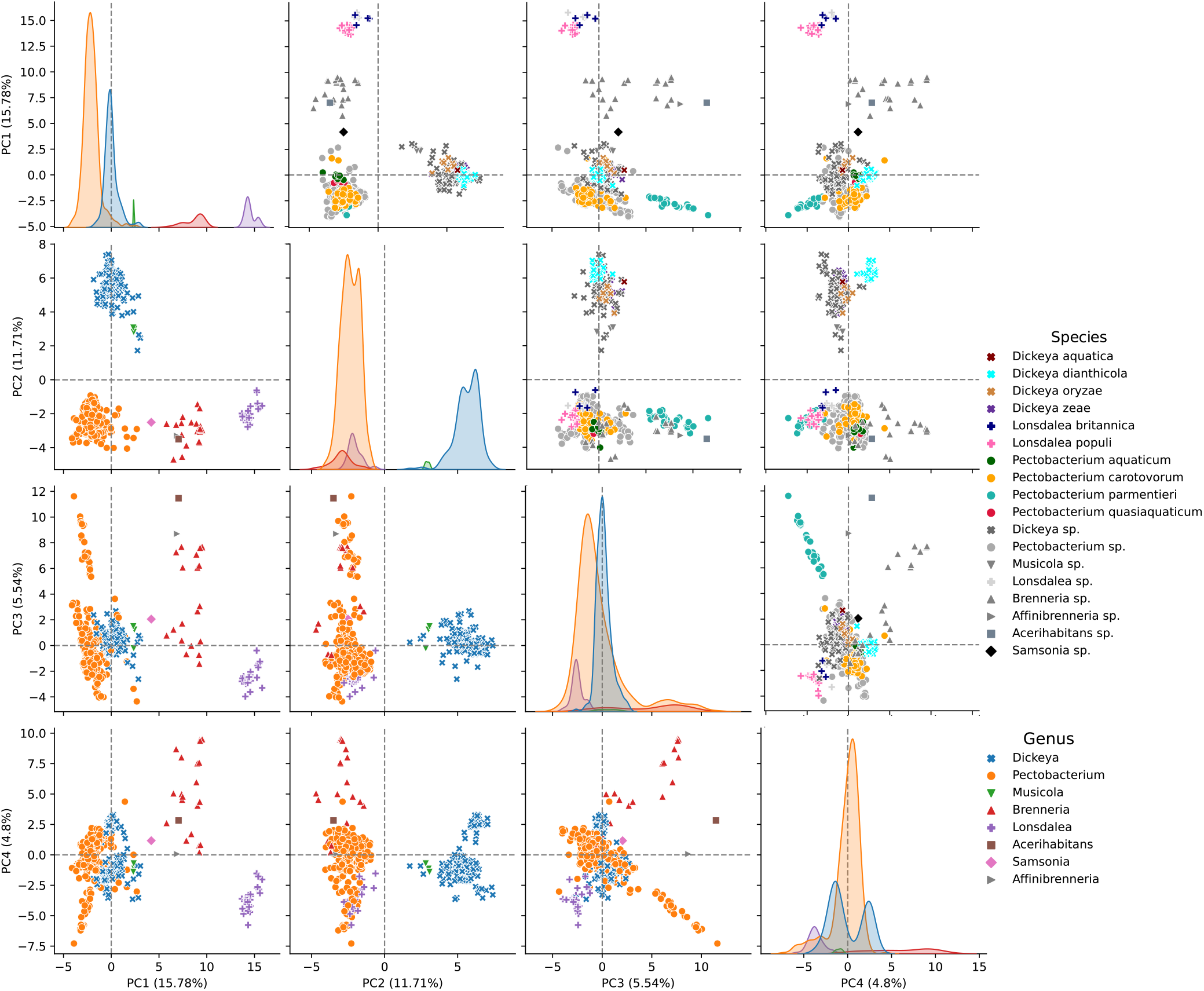
Principal component analysis of CAZy family frequencies in *Pectobacteriaceae*, projecting genomes onto all combinations of principal components (PCs) PC1-PC4. KDE plots (univariate distribution plots) are plotted on the diagonal, showing the marginal distribution of genomes in each column, colour coded by genus. Scatter plots below the KDE plots are colour coded and markers styled by genus classification, scatter plots above the KDE plots are styled by genus classification and selected species are colour coded by their species classification.

*Dickeya* and *Musicola* are the only genera to cluster in the positive PC2 axis, separately from *Pectobacterium* and all other genera (figure 2). *Pectobacterium* and *Dickeya* have plant hosts in common (66, 67), despite differences in their CAZome compositions. CAZome differences represented along the PC2 axis may therefore not be strongly related to host preference, but instead to lineage. However, the true (comprehensive) plant host range of *Pectobacterium* and *Dickeya* is unknown, so this may still be associated with host preference (68).

Notably, three *Dickeya poaceiphila* genomes cluster with *Musicola* genomes and away from the larger *Dickeya* cluster (SI figure 10). The genus *Musicola* was created from the reclassification of *Dickeya paradisiaca* (36), and analysis of the distance-based *Pectobacteriaceae* tree generated by ANIm (see Methods, figure 8) and a previously published phylogenetic analysis might support the assignment of these *Dickeya* genomes to the *Musicola* genus (36). A comparison of Biolog screens of 190 growing conditions retrieved from the literature (36, 69) indicate *Musicola paradisiaca, Musicola keenii*, and *D. poaceiphila* share a similar biochemical phenotype, differing on only 19 Biolog conditions (10%), seven of which are the ability to utilise particular sugars as a sole carbon source (D-mannitol, L-rhamnose, *α*-methyl-D-galactoside, *α*-D-lactose, pectin, D-arabinose, gentiobiose, *β*-methyl-D-galactoside, and D-tagatose) (SI table 18). These differences in sugar metabolism, which correlate with the similarity in CAZome composition implied by the clustering in 2, might form a basis for further taxonomic refinement.

Projection of genomes onto PC2, PC3 and PC4 does not cluster the genomes into distinct groups representing their respective genera, although the genomes do form clusters by species (figure 2). This suggests that PC1 and PC2 may capture most if not all genus-specific difference in CAZome composition for this group, and that PC3 and PC4 may represent diversity in *Pectobacteriaceae* resulting from more specific niche adaptation than preference for soft or hard plant tissue. For example, *P. aquaticum* and *P. quasiaquaticum* distinctly reside in fresh water and target aquatic flora, and cluster together but separately from other *Pectobacterium* spp. *L. populi* genomes, which uniquely among *Lonsdalea* causes bark cankers in poplar trees, forms a separate cluster to other members of the genus. Additionally, *Pectobacterium parmentieri* genomes form a distinct cluster when projecting genomes onto PC3 against PC4. *P. parmentieri* makes up a significant proportion of soft-rotting *Pectobacteriaceae* isolates that are found only in the temperate climates of the northern hemisphere, distinguishing it from *P. carotovorum* which has a global distribution and represents a significant fraction of *Pectobacteriaceae* isolates (70).

A loadings plot (figure 4) representing the degree of correlation between each CAZy family and projections along PCA axes PC1 and PC2 identifies several families that strongly correlate with a negative PC1 value (and thus potentially a preference for soft tissue degradation). Many of these are PL families, consistent with the PL expansion noted for SRP in figure 1. No PL families strongly correlate with a positive PC1 value (associated with hard plant tissue targeting *Pectobacteriaceae*). We interpret these data as support for the soft/hard tissue phenotype being associated with PL family expansion.

**Fig. 3.**
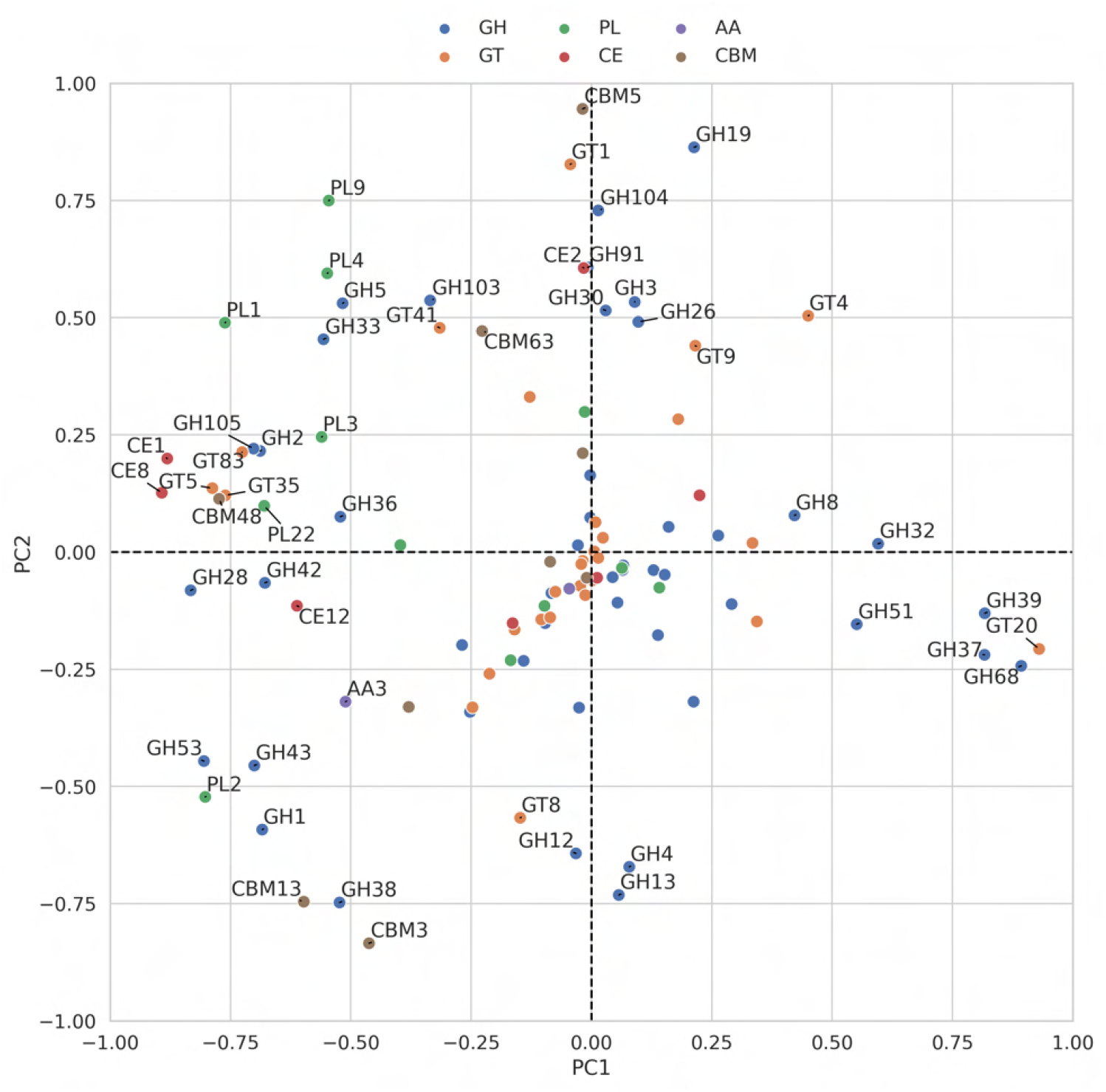
Loadings plot

**Fig. 4.** Loadings of CAZy families for PC1 against PC2 from principal component analysis of CAZy family frequencies in *Pectobacteriaceae*. CAZy families with a loading magnidtude of greater than 0.4 are annotated.

#### Pectobacteriaceae CAZy family saturation

To evaluate the completeness of the CAZome annotations (in terms of identifying all potential CAZy families) and the extent to which further CAZy family representatives remain to be discovered in *Pectobacteriaceae*, rarefaction plots were generated for all *Pectobacteriaceae* and for each genus separately (figure 5) (see Methods). All rarefaction curves are initially steep and most rapidly approach a plateau, suggesting CAZome saturation is close to being achieved with the current sampling effort and that the current dataset identifies most CAZy families present in the total *Pectobacteriaceae* CAZome. However, the curves do not become completely flat, so further sampling is likely to yield further new CAZy families. In particular the curves for *Musicola* and *Brenneria* do not plateau, suggesting that further sampling of these genera is likely to be the most productive in revealing additional *Pectobacteriaceae* CAZy families.

**Fig. 5.**
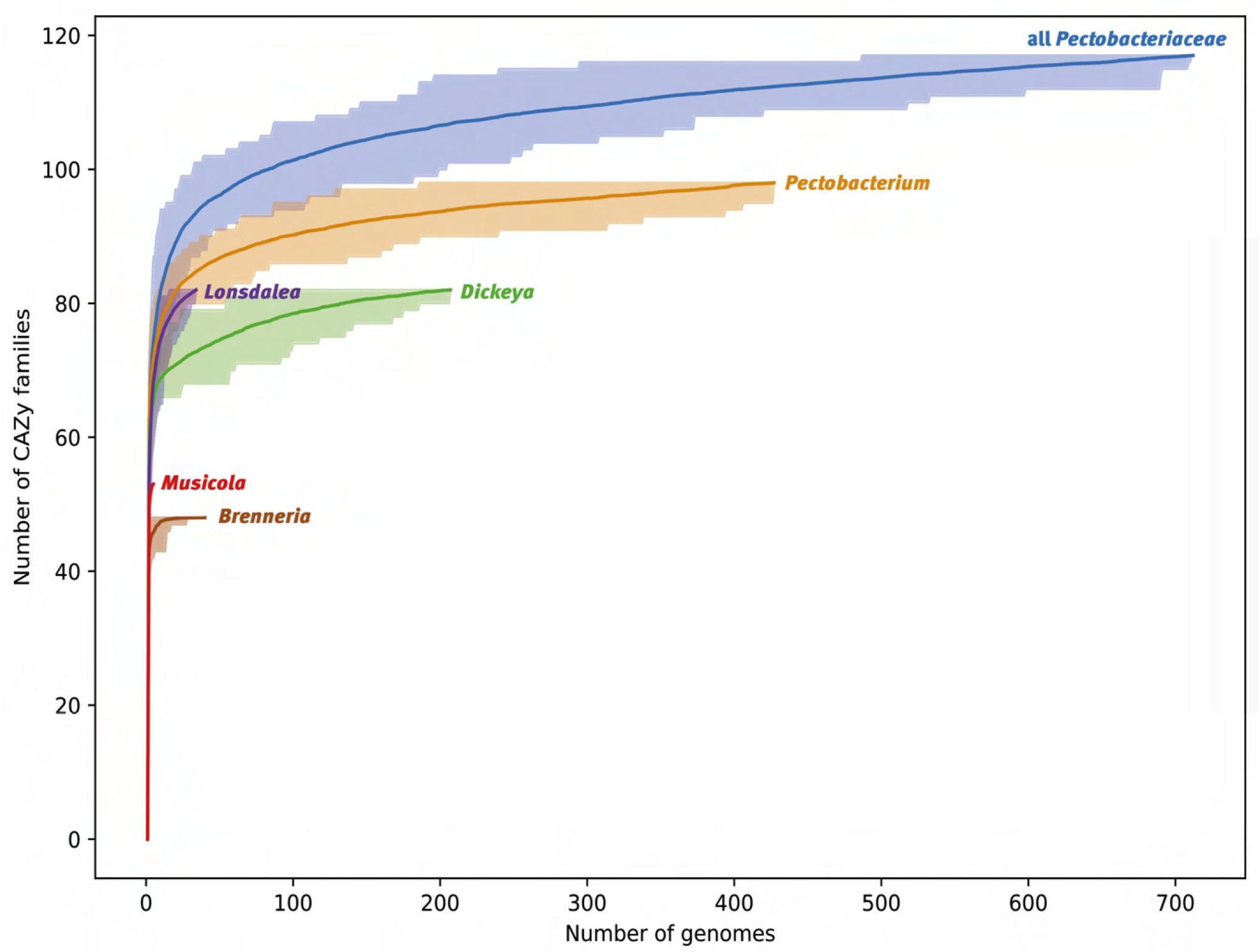
Rarefaction plots with the number of CAZy families observed per number of randomly sampled genomes. For each genus, 100 runs were performed and the variation in number of CAZyme families counted is shown by the ribbon area, while the average number of CAZy families across the runs is shown by the solid line. From the top lines represent all Pectobacteriaceae (blue), *Pectobacterium* spp. (tan), *Lonsdaleae* spp. (purple), *Dickeya* spp. (green), *Musicola* spp. (red), and *Brenneria* spp. (brown)

Figure 5 reflects the diversity of CAZome size and CAZy family frequency indicated by the PCA analysis in figure 2. Soft plant tissue targeting *Pectobacterium* and *Dickeya* possess more diverse CAZomes than the hard plant tissue targeting genera.

#### Differential expansion of CAZy families in soft and hard plant tissue targeting species

CAZy family frequencies were calculated and plotted as a clustermap (figure 6; see Methods). Clustering genomes by CAZyme family frequencies recapitulated genus classification, except for six *Pectobacterium* genomes not included within the main *Pectobacterium* sub-tree (figure 6[A]; SI table 19). These genomes were annotated with fewer CAZymes than other *Pectobacterium* genomes, possibly as the result of incomplete genomic sequences and/or annotation, consistent with a CheckM (71) analysis that reported 81-95% completeness with 2-9% contamination. Additionally, genome annotations for assembly GCA_029023745.1 (*Pectobacterium colocasium*) from the Prokaryotic Genome Annotation Pipeline (PGAP) output contained a high proportion (>30%) of frameshifted proteins, implying a low quality assembly resulting in annotation that underestimates CAZyme features (72). The annotated proteomes of these assemblies were excluded from subsequent analyses.

The clustering in figure 6 divides genomes with soft tissue preference from those with hard plant tissue targeting phenotypes, as two distinct clades. There is visible commonality in the pattern of CAZy family frequencies across genomes with the same SRP or HTD phenotype, although within this there are distinct genus-specific patterns of CAZy family representation (SI table 20). Four PL families (PL10, PL11, PL22 and PL26) are found only in soft plant tissue targeting genera, correlating with PL expansion in SRP. AA expansion in soft plant tissue targeting genera (figure 1) is reflected here, as expansion of the AA3 family.

**Fig. 6.**
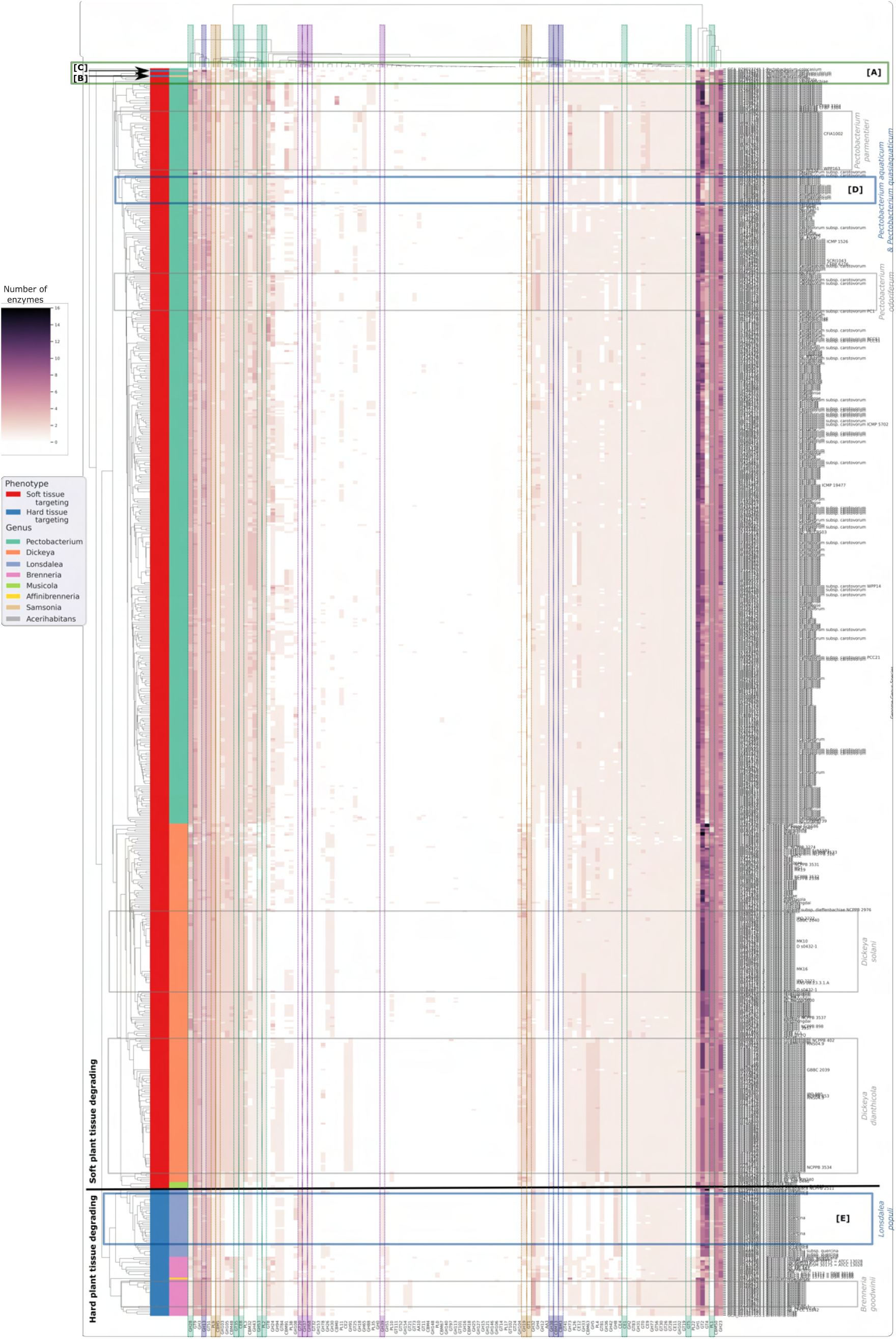
A clustermap of the frequency of CAZymes per CAZy family in *Pectobacteriaceae*. Rows represent genomes, columns represent CAZy families. Row marker colours indicate (left to right) the soft/hard plant tissue degrading and genus classification of each genomes. [A] six outlying *Pectobacterium* genomes, [B] outlying *Samsonia* genome, [C] outlying *Acerihabitans* genome that form a separate cluster from the other *Pectobacterium* genomes associated with low quality assembly. Genomes from the same species that are clustered together and have a distinct pattern of CAZy family frequencies are outlined in grey, including [D] *Pectobacterium aquaticum* and *Pectobacterium quasiaquaticum*, and [E] *Lonsdalea populi* which are outlined in blue.

CAZy families with a strong loading on PC1 (loading score magnitude greater than 0.75) in figure 4 are outlined in green (SRP) and blue (HTD) for their respective potential association with phenotype in figure 6. Families GH37, GH39 and GT20 were only found in hard plant tissue targeting genomes (SI table 20). Similarly, CAZy family GH68 is almost exclusive to hard plant tissue targeting genera, and is present in only four SRP genomes (figure 6).

No CAZy families that strongly associate with a soft plant tissue targeting phenotype in the PCA were found exclusively in SRP genomes, but families CE1, CE8, PL2, GH28 and GH53 are almost entirely absent from hard plant tissue targeting genomes (figure 6). Likewise, families GT5 and GT35 are present in nearly all soft plant tissue targeting genomes, but absent from all hard tissue targeting genomes except those of *Brenneria*. Additionally, CAZy family PL1 was found in both hard and soft plant tissue degrading genomes, but was more abundant - potentially indicating expansion within this functional group - in soft tissue degrading genomes.

CAZy families with a PC2 loading score magnitude greater than 0.7 (i.e. distinguishing *Dickeya* and *Musicola*) fromh the other genera) are shown in figure 6. Of these families, CBM13 is found exclusively in *Pectobacterium*, and families GH38, CBM3 and CBM3 are absent from *Dickeya* and *Musicola* (SI table 20). GH19 and GT1 are found in abundance in *Dickeya* but absent in most other genera. GH13, PL9 and CBM5 have extreme PC2 loadings and, while they are found in nearly all *Pectobacteriaceae* genomes, their abundance differentiates hard from soft tissue degrading genomes. We interpret these findings to suggest that a common set of CAZyme families is found across *Pectobaceriaceae* genomes, but their abundance and relative representation may vary by niche adaptation. Overlaid on this is a pattern of CAZy family presence and absence that is associated with host tissue preference and/or lineage.

#### Pectobacteriaceae contain a core CAZome of seven CAZyme families

CAZymes are essential to microbial growth and reproduction, so it seems likely that *Pectobacteriaceae* possess a core CAZome that is associated with core microbial processes and/or general plant tissue degradation.

In the absence of comprehensive functional characterisation we define the strict core CAZome as those CAZy families present in all 711 *Pectobacteriaceae* genomes.

Diversity of *Pectobacteriaceae* CAZome composition is such that only 7 of the 117 observed CAZy families (6%) in *Pectobacteriaceae* are present in all genomes, and constitute the strict core CAZome: GH3, GH23, GT2, GT9, GT51, CBM5 and CBM50. The mean frequency of these seven families is consistent across the genera (other than GT2 which appears to be more prevalent in *Acerihabitans* and *Affinibrenneria*, SI figure 11).

This core CAZome is predominantly made up of CAZy families GT2 (26% of CAZymes in the core CAZome), GH23 (21%) and CBM50 (19%). The consistently high frequency of GT2 and CBM50 CAZyme domains may not indicate functional redundancy, as GT2 domains target a variety of *β*-glucan linkages and CBM50 domains are known for glycan binding functions associated with several catalytic CAZy families. Both are associated with multiple biological processes, and individual family members may have differing specificities (73–75). GH23 CAZyme domains (involved in peptidoglycan lysis) and GT9 domains (heptosyltransferases) may be prominent in the core CAZome owing to their roles in the essential synthesis and maintenance of the protective bacteria outer membrane and its extracellular components (76– 78). The CBM5 domain is also typically not associated with plant pathogenic activity, being a chitin binding module that, along with an associated chitinase domain, is likely involved in degradation of chitinous material (79). These functional annotations suggest that the core CAZome of *Pectobacteriaceae* is probably not directly related to plant pathogenic activity.

#### Distinct groups of co-occurring CAZy families may be associated with soft and hard plant tissue targeting phenotypes

We further screened the 711 *Pectobacteriaceae* genome set for CAZy families that always co-occur in (i) any genus represented by more than one genome (excluding *Affinibrenneria, Acerihabitans* and *Samsonia*); (ii) all soft plant tissue targeting genomes (*Pectobacterium, Dickeya* and *Musicola*); and (iii) all hard plant tissue targeting genomes (*Lonsdalea, Brenneria, Affinibrenneria, Acerihabitans* and *Samsonia*) (figure 7) (see Methods).

**Fig. 7.**
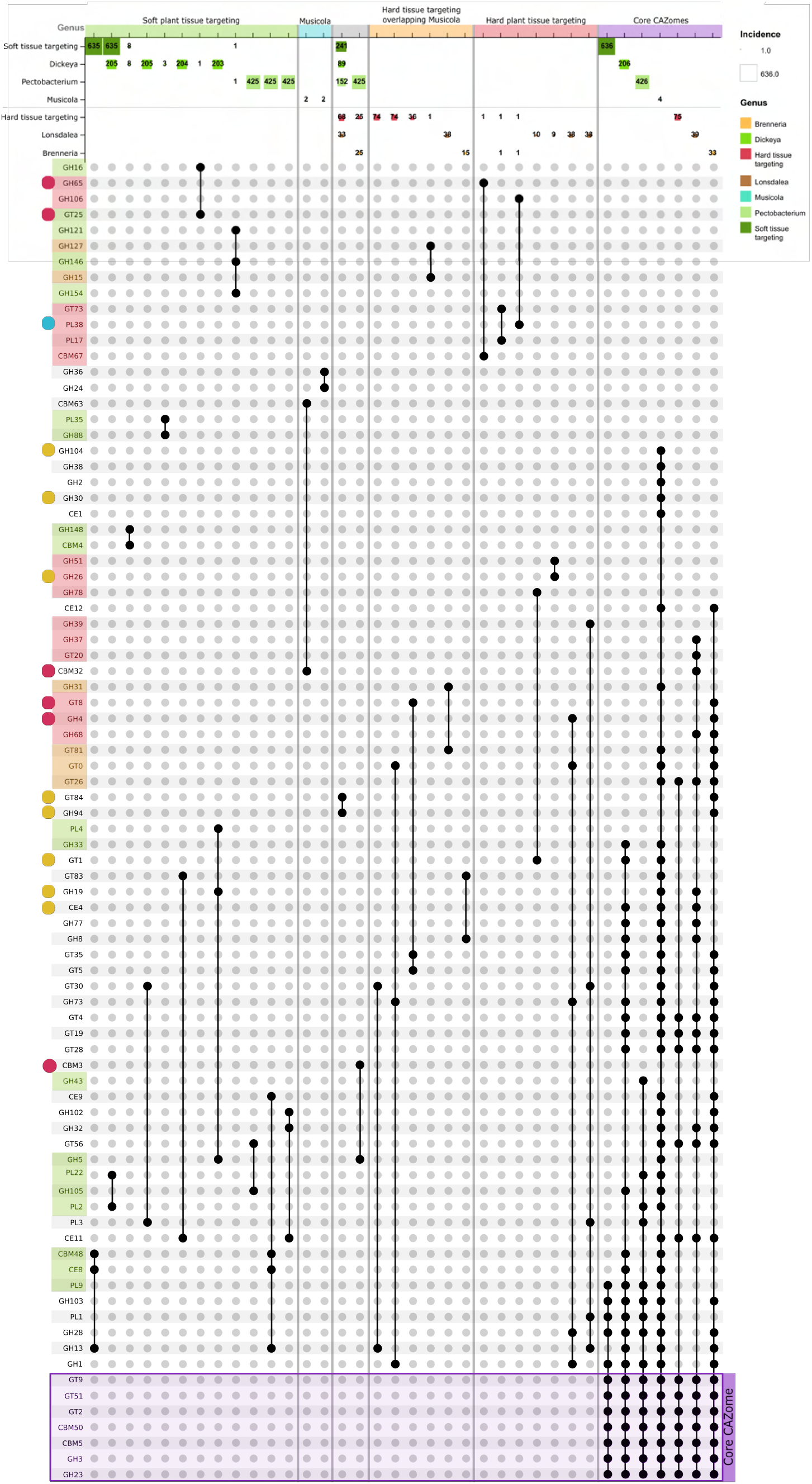
Upset plot of co-occurring CAZy families found across *Pectobacteriaceae*. Groups are collected from left to right by phenotype (soft plant tissue targeting genera *Pectobacterium, Dickeya* and *Musicola*, highlighted in green; hard plant tissue-targeting *Lonsdalea, Brenneria, Affinibrenneria, Acerihabitans*, and *Samsonia*, highlighted in pink). CAZy families present in all *Pectobacteriaceae* genomes (the core CAZome) are highlighted in purple. The names of families found to co-occur in only soft tissue degraders are highlighted in green; those found only in hard tissue degraders in pink; and those found only in hard tissue degraders plus *Musicola* in orange. Families annotated with the same colour circle (red, orange, blue; left of figure), were additionally indicated by `coinfinder` to co-occur together more often than expected when accounting for lineage.

A unique core CAZome was found for each genus. Taken together these illustrate distinct differences between soft and hard plant tissue targeting CAZomes (figure 7). Soft plant tissue-targeting *Pectobacterium, Dickeya* and *Musicola* genera extend the strict core *Pectobacteriaceae* CAZome to include families GH1, GH28, GH103, PL1 and PL9. Hard plant tissue targeting *Pectobacteriaceae* extend the strict core CAZome to include five glycosidic bond forming families: GT4, GT19, GT26, GT28, GT56, and family CE11. The *Musicola* core CAZome shows commonalities with those of both soft and hard tissue degrading genera.

We interpret these observations to indicate that selective pressure has applied not only to individual CAZyme functions (approximated by a single CAZy family) but to biological processes requiring co-ordinated activities of multiple CAZyme functions. For example, co-occurring CAZy families found only in soft- or hard tissue-degrading genomes may result from adaptation to host material requiring distinct sets of multiple CAZyme functions to act in concert to degrade host material.

#### Pectobacteriaceae genera and species contain co-evolving networks of co-associating CAZy families

The requirement for families always to co-occur within a group of organisms is a strict measure that will overlook CAZyme families that do not always occur together, but that still share correlated patterns of gain and loss, perhaps due to common contributions to a pathway or phenotype. We used `coinfinder`to analyse the occurrence of CAZy family pairs to determine whether there was evidence of a coincident relationship, i.e. one in which families were observed together in a CAZome more often than expected by chance, when corrected for lineage effects (48).

We used a distance-based tree reconstructed from ANIm analysis to estimate the lineage across *Pectobacteriaceae*, excluding all genomes with an assembly status of “contig” so that potentially incomplete sequences would not be of detriment to the analysis. Three pairs of redundant genomes were identified and only one genome from each included in the `coinfinder`to explore the association between CAZy families in the remaining 311 *Pectobacteriaceae* genomes. Networks of associating CAZy families found by `coinfinder`are presented in figure 8 and highlighted using circles where present in the co-occurring families clustermap (figure 7). An alternative presentation of the plot including taxonomic annotations is presented in SI figure 12.`coinfinder`identifies three co-evolving networks of CAZy families. These networks visibly partition the *Pectobacteriaceae* genomes by genus and species (figure 8).

**Fig. 8.**
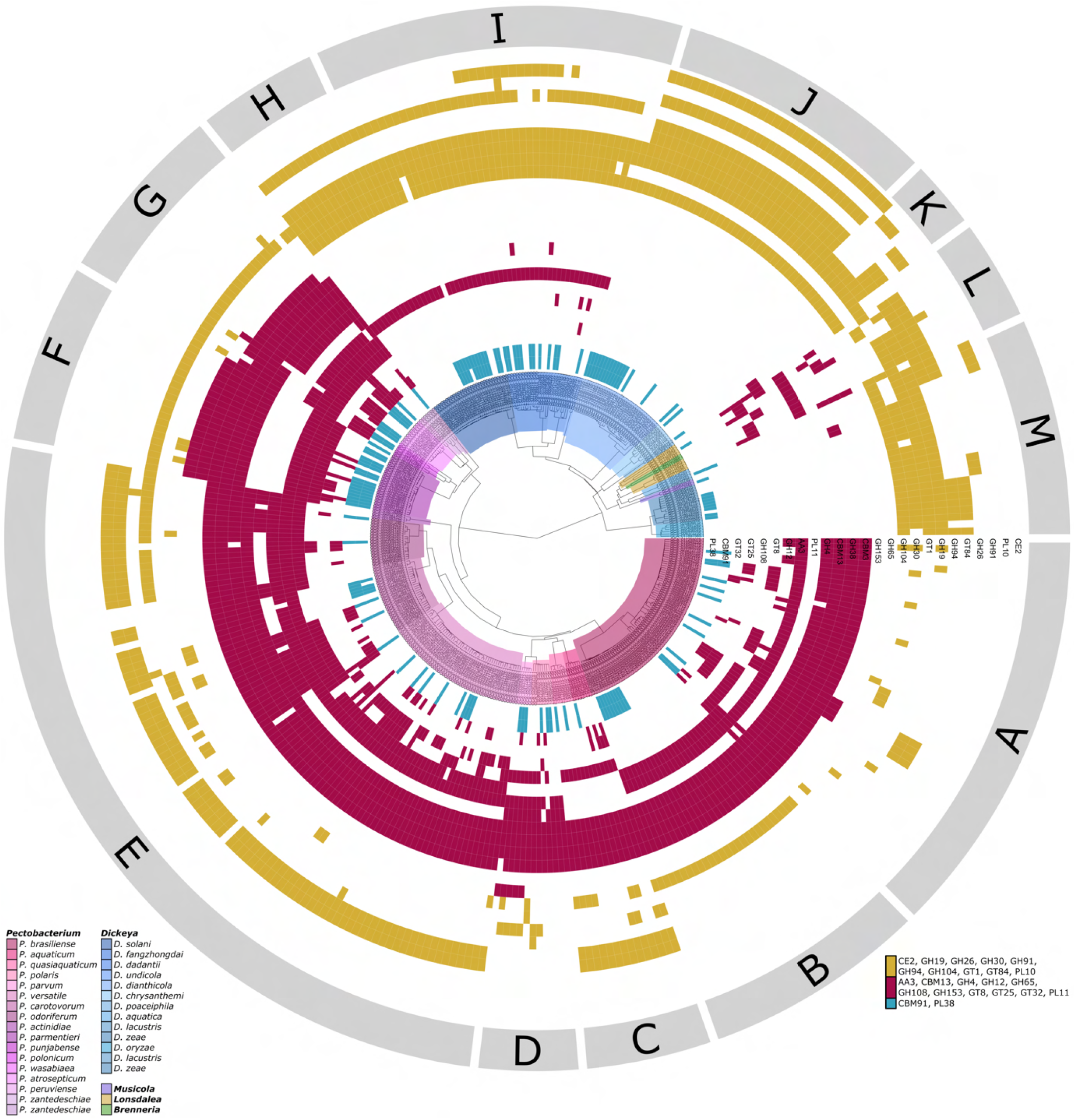
Output from `coinfinder`marking the presence and absence of co-evolving CAZy families in 311 Pectobacteriaceae genomes. The ANIm-derived dendrogram is colour coded by genus classification. CAZy families shaded the same colour (blue, red., yellow) were predicted to co-evolve together.

Two networks (maroon and gold) are primarily associated with *Pectobacterium* and *Dickeya*, respectively (figure 8). `coinfinder`did not identify a hard plant tissue targeting-specific co-evolving network. However, members of the *Pectobacterium* and *Dickeya*-associated networks are also found to co-occur in hard plant tissue degrading genera (group L in figure 8).

We subdivide genomes in figure 8 into thirteen groups (labelled A-M) on the basis of coevolving CAZyme family networks identified by `coinfinder`. Most species are placed in only one of these groups (see SI figure 12), but *P. brasiliense* is split between groups A and B, and *D. solani* genomes are split between groups H and I, with *D. fangzhongdai, D. dadantii*, and *D. undicola*.

The smallest network of co-evolving CAZy families identified by *coinfinder* comprises the pair of families CBM91 and PL38. This is the only network that does not visibly appear to be associated with a genus or phenotype, so might represent CAZyme families that participate one or processes that are not responsible for the characteristic phenotype or host range of any particular species.

The largest co-evolving network is found primarily in *Pectobacterium* (coloured maroon in figure 8). Families CBM13 and PL11 in this network are found exclusively in *Pectobacterium*, and family AA3 is only found in soft plant tissue targeting *Pectobacteriaceae* genomes (SI table 20).

The remaining network of co-evolving families is found predominantly in *Dickeya* (gold in figure 8). Families CE2, GH91 and PL10 from this network are only found in *Dickeya* (SI table 20). Other families in this network, such as GH94 and GT84 always co-occur together but are not unique to *Dickeya*.

Some families from the larger networks also associate together in *Brenneria, Lonsdalea* and *Acerihabitans*. Families GH4 and GT8 from the *Pectobacterium* network are also found in the *Brenneria* core CAZome, and families GH19, GH94, GT1 and GT84 are widely present in *Pectobacteriaceae* genomes.

`coinfinder`analysis indicates that there is evidence for a signal of mutual co-evolution and co-occurrence between CAZyme families that implies evolutionary pressure may apply to groups of enzymes working synergistically to complete a common biological process. This observation of lineage-associated co-evolving CAZy family networks is consistent with an association between niche adaptation and CAZome composition. Patterns of CAZy family associations that distinguish between pathogens with a prefence for soft or hard tissue suggests these processes are related to the phytopathogenic nature of these bacteria and their adaptation to particular plant hosts.

## Discussion

Industrial processing of globally abundant and renewable lignocellulose into bio-based products is a promising strategy for achieving carbon neutrality and a circular economy. The complexity of lignocellulose architecture necessitates the use of a large CAZyme arsenal with broad substrate specificities for its efficient degradation (21). However, the list of industrially exploited CAZymes is not exhaustive, and bioprocessing of lignocellulosic biomass is often inefficient compared to industry standards for other feedstocks (80). Data mining the CAZomes of phytopathogens, which constitute large CAZyme arsenals with broad substrate specificities that mirror the complexity and diversity in lignocellulosic architecture, is an existing strategy for identifying crude enzyme mixes and individual CAZymes for industrial exploitation of lignocellulose degradation with abundant potential for expansion and improvement (13, 21, 23).

We developed the software tool `cazomevolve`to facilitate bioinformatic screening of taxonomically and phenotypically diverse genome data sets to identify microbial CAZyme complements. By combining the annotation power of CAZy (via cazy_webscraper) and the CAZyme classifier dbCAN, `cazomevolve`automates the identification and analysis of comprehensive CAZome data sets in microbial genomes. This functionality complements the alternative SACCHARIS pipeline that streamlines detection of new CAZyme or CBM specificities in CAZyme families of interest (81). SACCHARIS explores diversity within CAZyme families, reconstructing gene trees to identify under-explored or underrepresented sequences (81). By contrast `cazomevolve`explores the entire CAZyme complement of the input genomes, augmenting existing annotations and using unsupervised machine learning to identify trends between CAZyme families, phylogeny and phenotype.

Using cazomevolve we extended CAZyme annotation in *Pectobacteriaceae* from 17,000 canonical representatives in the CAZy database to over 78,000 CAZymes. This enabled identification of associations between CAZome composition, niche, phenotype, and taxonomy, as well as several co-evolving groups of CAZyme families that may be associated with host preference and which could potentially form the basis of novel industrial enzyme mixed. Specifically, we found: (i) differential expansion of polysaccharide lyase (PL) and auxiliary activity (AA) CAZymes in soft plant tissue targeting genomes; (ii) differential expansion of CAZyme families in species associated with distinct niches and plant host ranges; (iii) distinct CAZome compositions in soft and hard plant tissue targeting genomes; and (iv) lineage and phenotype-specific groups of co-evolving CAZyme families. Significant CAZome diversity between *Pectobacterium, Dickeya* and *Musicola* was also identified, despite these genera sharing overlapping plant host ranges.

Our analyses of the *Pectobacteriaceae* suggest that a wealth of novel CAZyme functions and synergistic associations between CAZyme families reamain to be discovered. The ability to automatically interrogate and screen taxonomically and phenotypically diverse CAZomes from plant-degrading microorganisms *in silico* will be invaluable for surveying and exploiting this functional landscape. The ability to infer reliably associations between CAZome composition, niche, phenotype, and taxonomy will facilitate prediction of host range from genome sequence. This capability may enable prediction of host range, host-jump potential, and the commercial and agricultural impact of emerging phytopathogens, informing strategies to limit the impact of emerging phytopathogens and improve food security.

### Diversity of Pectobacteriaceae CAZomes correlates with soft and hard plant tissue targeting phenotypes

Lignocellulose composition differs between soft and hard plant tissues. Pectins are an extremely diverse class of polysaccharides that found more abundantly in soft plant tissues and soft tissues of new shoots owing to their role in controlling cell flexibility, and thus cell proliferation and plant growth (82, 83). Degradation of pectins is primarily performed by members of the polysaccharide lyase (PL) CAZy class (31). Therefore, it may be expected that soft plant tissue targeting species may have more PLs than hard PTD genera. We find expansion of PLs in soft plant tissue targeting *Pectobacterium* and *Dickeya* genera, relative to hard tissue targeting *Lonsdalea, Brenneria, Affinbrenneria*, and *Acerhabitans* (figure 1). This expansion of PLs correlates with an increased number of PL families (figure 6), with seven PL families found only in soft tissue targeting genomes, and with PL families that associated with soft plant tissue targeting species (i.e. had a strongly negative PC1 value). We found no PL families that strongly associated with hard plant tissue targeting species. These data indicate differential expansion of PLs in *Pectobacterium* and *Dickeya*, consistent with their soft plant tissue targeting phenotype, implying a more general association between niche adaptation, carbohydrate processing capabilities and CAZome composition in *Pectobacteriaceae*. SRP are opportunistic phytopathogens that reside in soil, waterways, and on plants, entering through openings (such as stoma and wounds), and targeting the soft tissues of food-crops (84, 85). Commonality of CAZome composition in *Pectobacterium* and *Dickeya* may reflect evolution of a general-purpose soft plant tissue degrading enzyme mix enabling these bacteria to target an overlapping range of plant species. Such a mixture could be a valuable addition to the arsenal of industrial biotechnology tools in this sector.

As *Musicola* is represented by only four genomes in this analysis, while *Samsonia, Acerhabitans*, and *Affinibrenneria* genera are each represented by a single genome, there is insufficient data to assess confidently variation within these genera, or to assess if the genomes included in this analysis are representative of their respective genus. To ensure robustness, and potentially to identify novel CAZyme relationships, these analyses will need to be reapplied to this group as new genome assemblies become available, especially for underrepresented genera.

A CAZyme family may encompass multiple biochemical activities (86), therefore differences in precise carbon source specificity may not be detectable in the variation in CAZyme family frequencies at the level of our analyses. These more subtle differences in carbohydrate processing phenotypes might be more detectable when comparing EC numbers between each CAZome (87). However, precise functional characterisation of CAZymes is not comprehensive, and much practical cataloguing and characterisation work requires to be done for this to be a very useful prospect.

### Analysis of CAZome composition may facilitate prediction of host range and host jump potential

The association between CAZome composition and the ability to degrade specific plant host material may facilitate prediction of host range from genome sequence, or determination that an organism will *not* degrade a particular hose. This might enable prediction of host-jump potential and commercial impact of emerging phytopathogens, and enable genome-informed regulation and screening of phytopathogens to improve food security.

However, inferences of association between plant host range and geographical distribution with CAZome composition are restricted by our limited current understanding of the true or comprehensive plant host range of *Pectobacteriaceae* phytopathogens (88). A plant host is often only reported when the host is of economic interest (i.e. a food crop), but a wider host range may exist in the wild for such phytopathogens. Mapping of comprehensive host ranges onto CAZome compositions will require practical survey of plant polysaccharide composition as well as microbial CAZyme composition.

Nonetheless strong divergence in CAZome composition between soft and hard plant tissue targeting species, and distinct CAZome compositions in species with unique niches, suggests that these inferences may be possible.

### CAZome composition may facilitate taxonomic circumscription

The strong correlation between taxonomy, phenotype, and CAZome composition indicated by our analysis suggests that analysis of the CAZome may aid with circumscription of taxonomic groups (e.g. species and genus delineation). Trees based on CAZyme family frequencies showed similar topology to dendrograms generated using ANI analyis, which has been frequently used to assist taxonomic classification of *Pectobacteriaceae* genomes (36) (SI figure 13). This is consistent with the reported correlation between taxonomy and the topology of a dendrogram constructed from a CAZyme family presence/absence distance matrix in fungi (30). Moreover, estimating genome distances from distances between CAZome compositions appeared insensitive to incomplete genomic sequences, and relates directly to biochemical capability and phenotype, where simple sequence distance measures do not. Such analyses may add more refinement to automated phenotype prediction and taxonomic classification.

### Systematic identification of enzymes mixes for industrial exploitation

`cazomevolve`screens genomic sequences for CAZy families that co-occur, and maps CAZyme activities (represented by CAZy families) to taxonomy, phenotype, and plant host range. This systematically identifies groups of CAZymes that may have co-evolved to target specific lignocellulosic biomass compositions.

We identified three networks of co-evolving CAZy families, two of which are associated with specific *Pectobacteriaceae* lineages, which we interpret as a correlated adaptation of lignocellulosic degrading enzymes with potential synergistic activity in lignocellulose degradation. Identifying such groups of co-evolving synergistic CAZy families using `cazomevolve`and coinfinder, especially in conjunction with specific plant material targets, would form a basis for development of enzyme mixes for efficient degradation of a specific substrate’s lignocellulosic biomass.

## Conclusions

`cazomevolve`automates annotation and exploration of the entire CAZyme complement (the CAZome) of user defined genome datasets in a systematic and reproducible manner. By using `cazomevolve`to explore the taxonomically and phenotypically diverse *Pectobacteriaceae* family, we identified CAZyme sets that may be responsible for the distinct soft and hard plant tissue targeting phenotypic divide between *Pectobacteriaceae* genera. We found a strong association between taxonomy, plant host range, and CAZome composition, suggesting a role for CAZome composition in niche adaptation. Mapping of CAZyme activities to taxonomy and plant host range offers potential for identifying relationships between synergistic CAZyme activities and specific lignocellulosic architectures, and insight into design of enzyme mixes for broad and targeted degradation of lignocellulosic biomass, thus improving energy security. Identifying these associations may facilitate prediction of plant host range and host jump potential from genomic sequence alone, and be highly informative for devising strategies to limit the impact of existing and emerging phytopathogens, with positive impacts on food security and sustainability.

## Abbreviations

AA: Auxiliary activity
CBM: Carbohydrate binding module
CE: Carbohydrate esterase
EC: Enzyme Commission
GH: Glycoside hydrolase
GT: Glycosyl transferase
NCBI: National Centre for Biotechnology Information
PL: Polysaccharide lyases.

## Availability of data and materials

`cazomevolve`. Project name: `cazomevolve`

Project home page: https://hobnobmancer.github.io/cazomevolve

GitHub Repository: https://github.com/HobnobMancer/cazomevolve.

Documentation: https://cazomevolve.readthedocs.io/en/latest.

Operating systems: Linux, MacOS and Windows 7 or higher.

Programming language: Python 3.9 or higher. License: MIT License.

Any restrictions to use by non-academics: Commercial rights reserved.

### Supplementary data

Project name: SI Hobbs et al 2024 - Pectobacteriaceae

Project home page: https://hobnobmancer.github.io/SI_Hobbs_et_al_2024_Pecto/

GitHub Repository: https://github.com/HobnobMancer/SI_Hobbs_et_al_2024_Pecto.

Operating systems: Linux, MacOS and Windows 7 or higher. Programming language: Python 3.9.

License: MIT License.

Any restrictions to use by non-academics: Commercial rights reserved.

## Authors’ contributions

Conceptualization, E.E.M.H., T.M.G., L.P.; methodology, E.E.M.H., T.M.G. and L.P.; software, E.E.M.H.; validation, E.E.M.H., T.M.G. and L.P.; formal analysis, E.E.M.H., T.M.G. and L.P.; investigation, E.E.M.H., T.M.G. and L.P.; resources, E.E.M.H.; data curation, E.E.M.H.; writing— original draft preparation, E.E.M.H., T.M.G. and L.P.; writing—review and editing, E.E.M.H., T.M.G. and L.P.; visualization, E.E.M.H., T.M.G. and L.P.; supervision, T.M.G. and L.P.; project administration, T.M.G. and L.P.; funding acquisition, T.M.G. and L.P.. All authors have read and agreed to the published version of the manuscript.

## Consent for publication

Not applicable.

## Ethics approval and consent to participate

Not applicable.

## Additional files

Additional data files are available in the online repository, including the original (full size, high resolution) figures (https://github.com/HobnobMancer/SI_Hobbs_et_al_2024_Pecto).

Supplementary material is available in supplementary file ‘Supplementary Information for: Carbohydrate active enzymes in Pectobacteriaceae: coevolving enzyme sets and host adaptation’ (PDF, 2976KB, https://github.com/HobnobMancer/SI_Hobbs_et_al_2024_Pecto/blob/master/Hobbs_et_al_SI_Pectobacteriaceae.pdf).

