## Supplementary Information for "Carbohydrate active enzymes in *Pectobacteriaceae*: coevolving enzyme sets and host adaptation"

Emma E. M. Hobbs, Tracey M. Gloster, Leighton Pritchard

2026

This document contains all supplementary information for the Manuscript: Carbohydrate active enzymes in *Pectobacteriaceae*: coevolving enzyme sets and host adaptation.

#### Contents

**1 Methods** 2

**2 Comparison of proteome size** 6

**3 Comparison of the percentage of the proteome that is encapsulated in the CAZome** 8

**4 Comparison of CAZome size** 11

**5 Comparison of the number of CAZyme families** 14

**6 Comparison of CAZy class frequencies** 17

**7 Principal Component Analysis (PCA) of CAZy family frequencies** 27

**8 Exploration of CAZy family frequencies** 31

**9 Coinfinder output** 33

### 1 Methods

#### SI Figure 1: Method to compile a proteome dataset

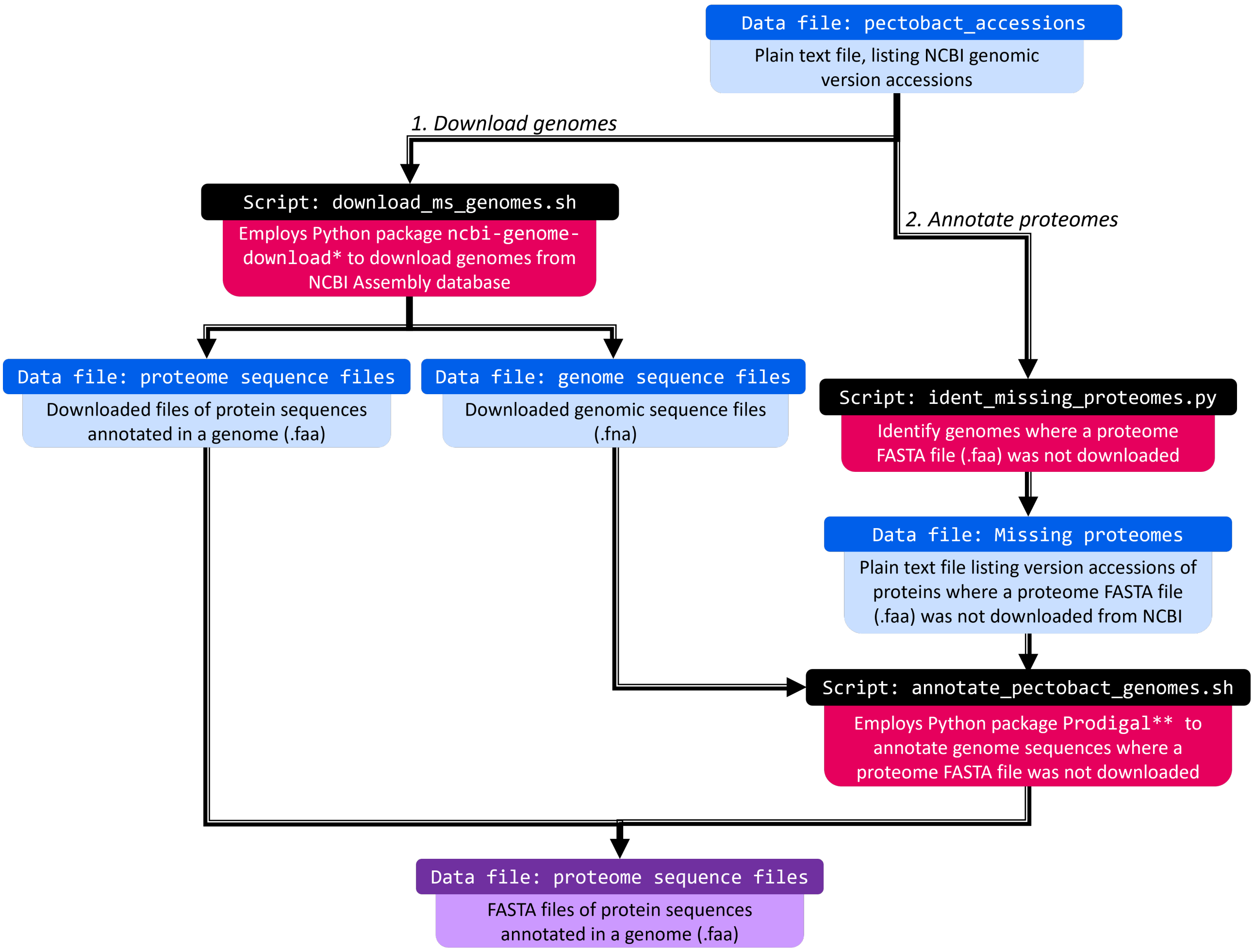


Figure 1: **Schematic summarising the method used to compile a proteome dataset for *Pectobacteriaceae*.** Bash scripts used to coordinate downloading genomes from NCBI (* using `ncbi-genome-download`), and annotating the downloaded genomes (** using `Prodigal`) to generate FASTA files of annotated protein sequences are shown in black. The final output is shaded in purple.

#### SI Figure 2: Method to compile a CAZyme dataset

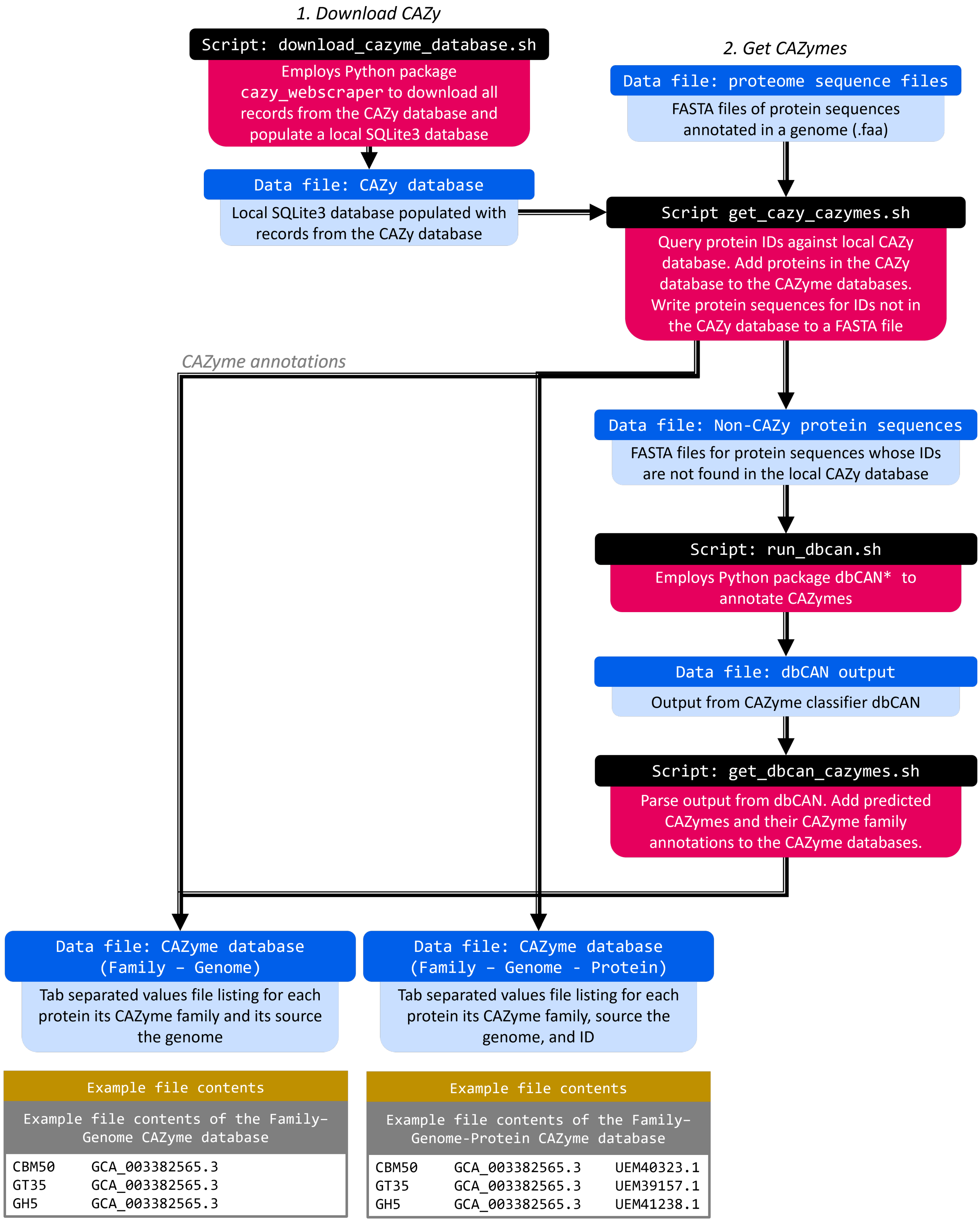


Figure 2: **Schematic summarising the method used to compile a *Pectobacteriaceae* CAZyme database**. Bash scripts (represented by black and pink boxes) used to coordinate `cazomevolve` to compile CAZyme databases listing CAZyme annotations from a local CAZy database, and predicted by dbCAN. Data files generated by the pipeline are represented by blue boxes. Example output are presented in grey boxes.

#### SI Figure 3: Method to add taxonomic classifications

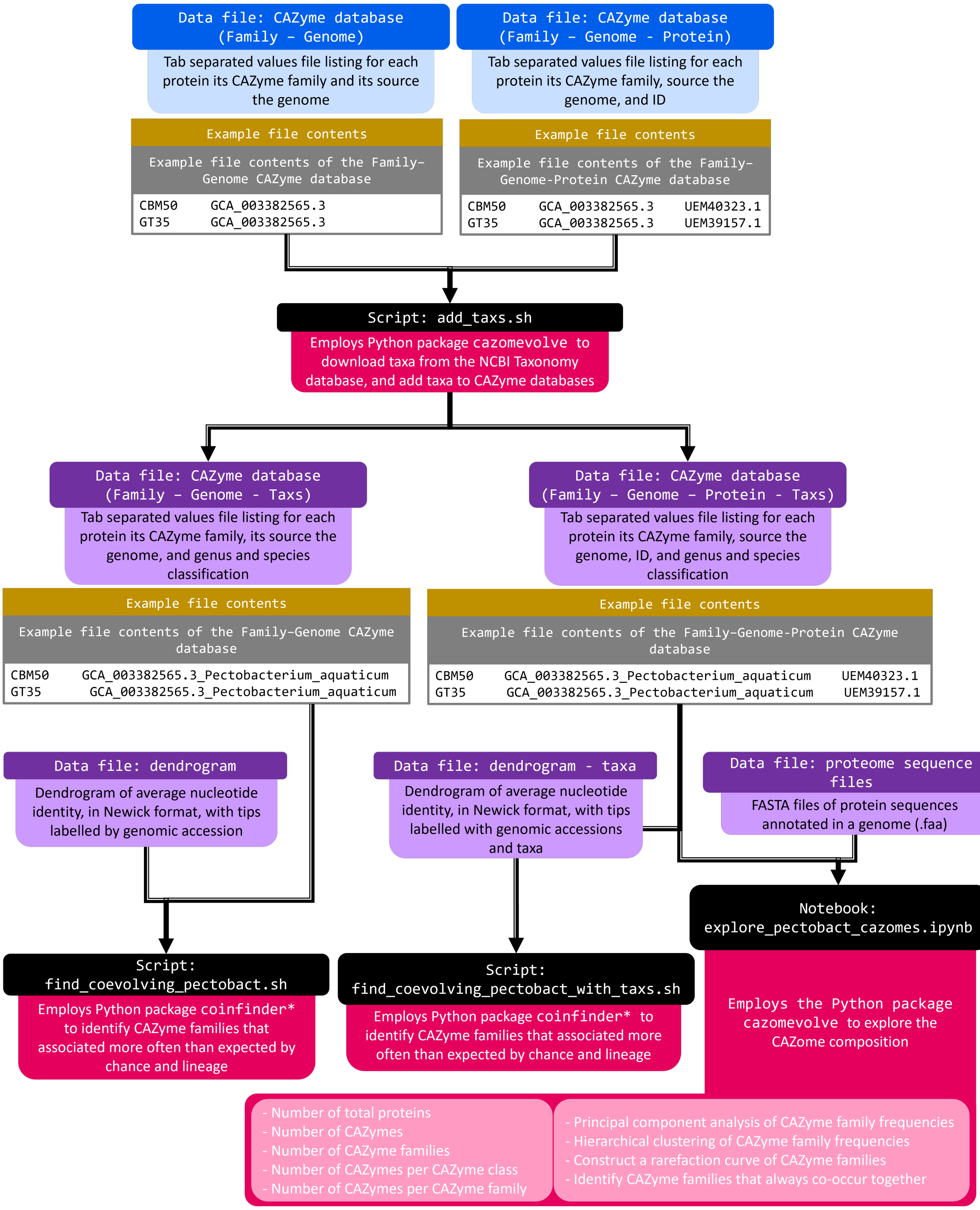


Figure 3: **Schematic summarising the methods used to add taxonomy data to the *Pectobacteriaceae* CAZyme databases** Scripts (represented by black and pink boxes) used to coordinate `cazomevolve` to download taxa (genus and species) from NCBI and add the taxa to CAZyme databases, as well as the application of the CAZyme databases within the `cazomevolve` pipeline. Operations in the pale pink boxes are performed by `cazomevolve`, configured using the notebook `explore_pectobact_cazomes.ipynb`. Blue boxes represent input datafiles. Purple boxes represent datafiles that were interrogated to compare the CAZome compositions of Pectobacteriaceae. Example content of datafiles is shown in grey boxes.

#### SI Figure 4: Method to build average nucleotide identity (ANI) tree

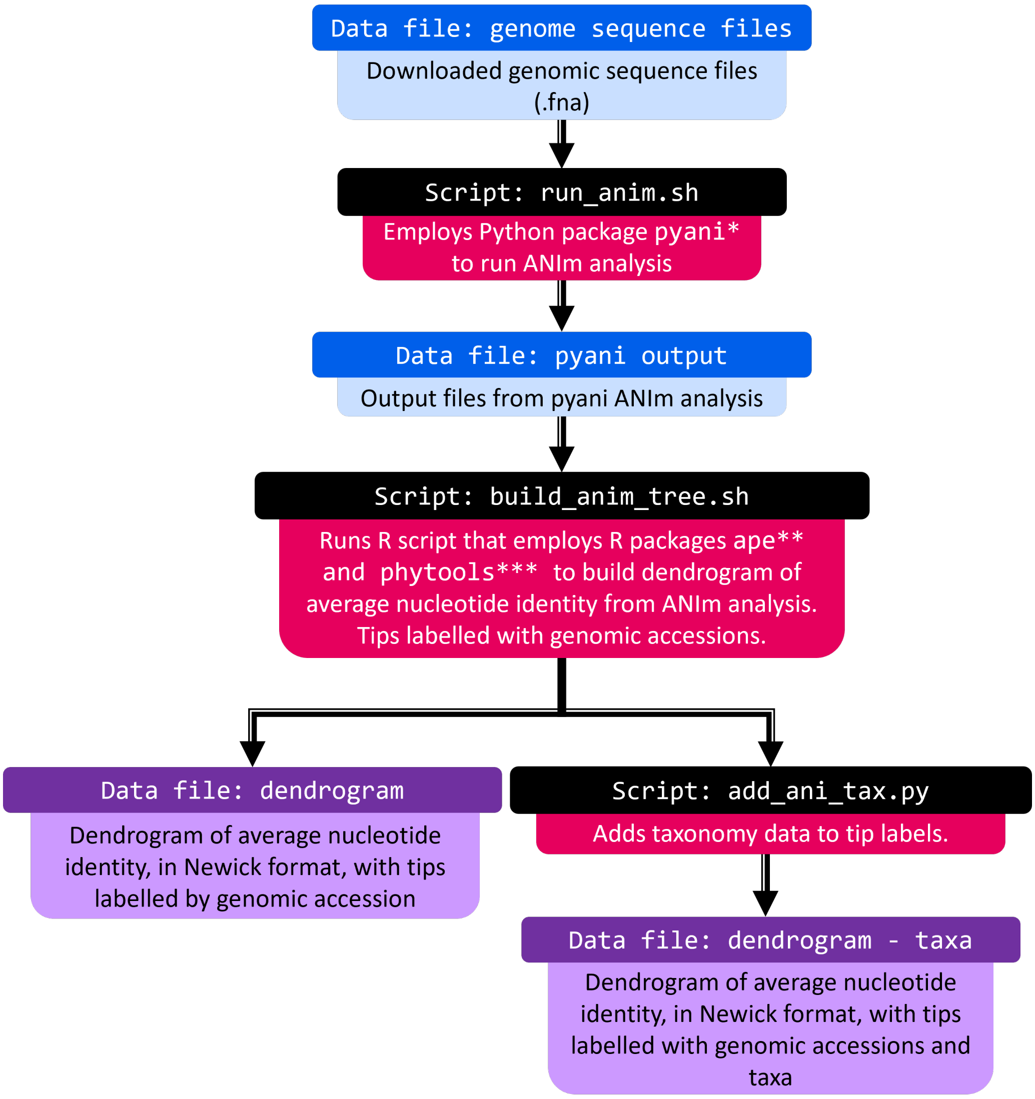


Figure 4: **Schematic summarising the method used to build a distance-based tree for *Pectobacteriaceae* genome sequences** Scripts used to coordinate `pyani` to calculate the average nucleotide identity, and using the R script to build a dendrogram.

#### 2 Comparison of proteome size

##### SI Figure 5: Boxplot overlaid by a scatter plot of the number of proteins in the proteome

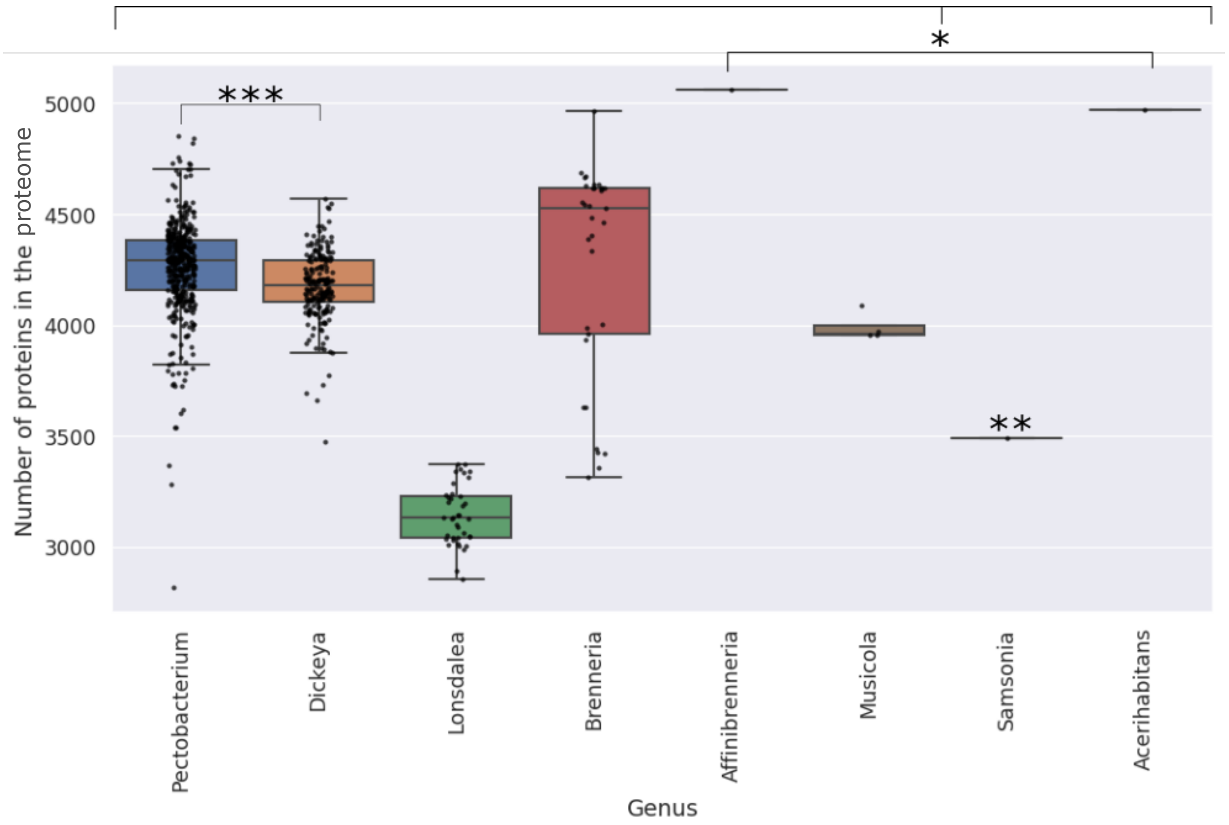


Figure 5: Number of proteins (i.e. the number of unique protein IDs) in each genome, grouped by genus with each point representing a unique genome, overlaying genomes are indicated with a darker shade.

#### SI Table 1: Tukey HDS post hoc test to identify genera with statistically significantly different mean proteome sizes

Output from a Tukey HDS post hoc test of the proteome sizes across the *Pectobacteriaceae* genera, performed using the Python StatsModels package (see Methods). A p-value (P-Adj, adjusted p-value) of 0.05 was used as a cut off rejecting the null hypothesis (NH). In addition the mean difference (MeanDiff) and lower and upper bounds of the confidence interval for the mean difference.

| Group1 | Group2 | MeanDiff | P-Adj | Lower | Upper | Reject NH |
|---|---|---|---|---|---|---|
| Acerihabitans | Affinibrenneria | 95 | 1 | -839.9267 | 1029.9267 | False |
| Acerihabitans | Brenneria | -698.7576 | 0.0344 | -1369.7924 | -27.7227 | True |
| Acerihabitans | Dickeya | -792.1408 | 0.0072 | -1454.8365 | -129.4451 | True |
| Acerihabitans | Lonsdalea | -1826.7179 | 0 | -2496.2329 | -1157.203 | True |
| Acerihabitans | Musicola | -977 | 0.0017 | -1716.1245 | -237.8755 | True |
| Acerihabitans | Pectobacterium | -708.287 | 0.0262 | -1370.1448 | -46.4293 | True |
| Acerihabitans | Samsonia | -1480 | 0 | -2414.9267 | -545.0733 | True |
| Affinibrenneria | Brenneria | -793.7576 | 0.0083 | -1464.7924 | -122.7227 | True |
| Affinibrenneria | Dickeya | -887.1408 | 0.0014 | -1549.8365 | -224.4451 | True |
| Affinibrenneria | Lonsdalea | -1921.7179 | 0 | -2591.2329 | -1252.203 | True |
| Affinibrenneria | Musicola | -1072 | 0.0003 | -1811.1245 | -332.8755 | True |
| Affinibrenneria | Pectobacterium | -803.287 | 0.0059 | -1465.1448 | -141.4293 | True |
| Affinibrenneria | Samsonia | -1575 | 0 | -2509.9267 | -640.0733 | True |
| Brenneria | Dickeya | -93.3832 | 0.3004 | -217.3402 | 30.5738 | False |
| Brenneria | Lonsdalea | -1127.9604 | 0 | -1284.3254 | -971.5954 | True |
| Brenneria | Musicola | -278.2424 | 0.2348 | -628.2492 | 71.7644 | False |
| Brenneria | Pectobacterium | -9.5295 | 1 | -128.9256 | 109.8667 | False |
| Brenneria | Samsonia | -781.2424 | 0.0101 | -1452.2773 | -110.2076 | True |
| Dickeya | Lonsdalea | -1034.5772 | 0 | -1150.0234 | -919.131 | True |
| Dickeya | Musicola | -184.8592 | 0.6978 | -518.5995 | 148.881 | False |
| Dickeya | Pectobacterium | 83.8537 | 0.0002 | 27.8783 | 139.8292 | True |
| Dickeya | Samsonia | -687.8592 | 0.0355 | -1350.5549 | -25.1635 | True |
| Lonsdalea | Musicola | 849.7179 | 0 | 502.634 | 1196.8019 | True |
| Lonsdalea | Pectobacterium | 1118.4309 | 0 | 1007.8962 | 1228.9657 | True |
| Lonsdalea | Samsonia | 346.7179 | 0.7657 | -322.797 | 1016.2329 | False |
| Musicola | Pectobacterium | 268.713 | 0.2146 | -63.3603 | 600.7863 | False |
| Musicola | Samsonia | -503 | 0.4363 | -1242.1245 | 236.1245 | False |
| Pectobacterium | Samsonia | -771.713 | 0.0099 | -1433.5707 | -109.8552 | True |

### 3 Comparison of the percentage of the proteome that is encapsulated in the CAZome

**SI Figure 6: Boxplot overlaid by a scatter plot of the percentage of the proteome encapsulated by the CAZome**

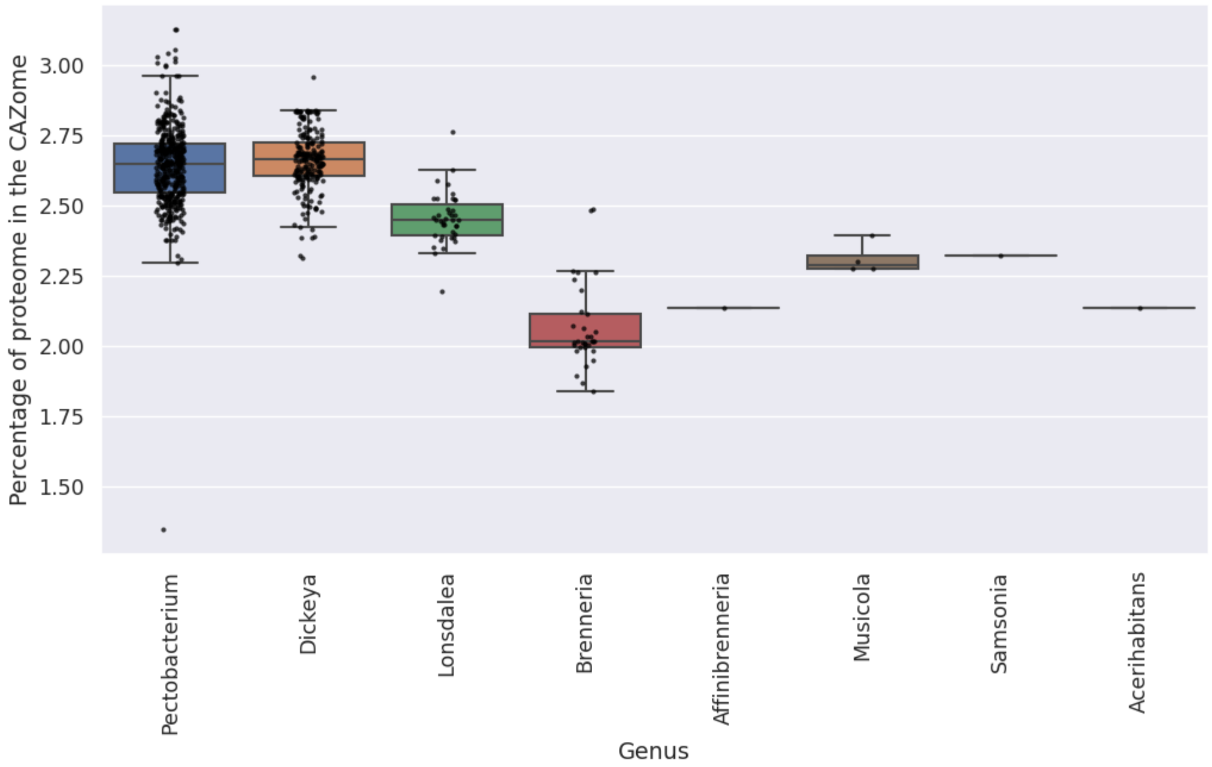


Figure 6: The percentage of the proteome encapsulated by the CAZome in each genome, grouped by genus with each point representing a unique genome, overlaying genomes are indicated with a darker shade.

#### SI Table 2: Tukey HDS post hoc test to identify statistically significant differences in mean percentage of the proteome that is encapsulated by the CAZome in soft and hard plant tissue targeting genomes

Output from a Tukey HDS post hoc test of the percentage of the proteome that is encapsulated by the CAZome across soft plant tissue targeting genera (*Pectobacterium* and *Dickeya*) and hard plant tissue targeting genera (*Acerhabitans*, *Affinibrenneria*, *Brenneria*, *Lonsdalea*, and *Samsonia*). *Musicola* is labelled as 'unknown' owing to its preference of soft and hard plant tissues not having yet been established in the literature. A p-value (P-Adj, adjusted p-value) of 0.05 was used as a cut off rejecting the null hypothesis (NH). In addition the mean difference (MeanDiff) and lower and upper bounds of the confidence interval for the mean difference.

| Group1 | Group2 | MeanDiff | P-Adj | Lower | Upper | Reject NH |
|---|---|---|---|---|---|---|
| Hard | Soft | 0.3753 | 0 | 0.3331 | 0.4175 | True |
| Hard | Unknown | 0.0371 | 0.8757 | -0.1404 | 0.2145 | False |
| Soft | Unknown | -0.3382 | 0 | -0.5117 | -0.1648 | True |

#### SI Table 3: Tukey HDS post hoc test to identify genera with statistically significantly different mean percentage of the proteome encapsulated in the CAZome

Output from a Tukey HDS post hoc test of the percentage of the proteome represented by the CAZome across the *Pectobacteriaceae* genera, performed using the Python StatsModels package (see Methods). A p-value (P-Adj, adjusted p-value) of 0.05 was used as a cut off rejecting the null hypothesis (NH). In addition the mean difference (MeanDiff) and lower and upper bounds of the confidence interval for the mean difference.

| Group1 | Group2 | MeanDiff | P-Adj | Lower | Upper | Reject NH |
|---|---|---|---|---|---|---|
| Acerihabitans | Affinibrenneria | -0.0005 | 1 | -0.5762 | 0.5751 | False |
| Acerihabitans | Brenneria | -0.0662 | 0.9997 | -0.4793 | 0.347 | False |
| Acerihabitans | Dickeya | 0.5273 | 0.0024 | 0.1192 | 0.9353 | True |
| Acerihabitans | Lonsdalea | 0.3215 | 0.2572 | -0.0907 | 0.7337 | False |
| Acerihabitans | Musicola | 0.1777 | 0.9356 | -0.2774 | 0.6327 | False |
| Acerihabitans | Pectobacterium | 0.5105 | 0.0038 | 0.103 | 0.918 | True |
| Acerihabitans | Samsonia | 0.1884 | 0.9752 | -0.3873 | 0.764 | False |
| Affinibrenneria | Brenneria | -0.0656 | 0.9997 | -0.4788 | 0.3475 | False |
| Affinibrenneria | Dickeya | 0.5278 | 0.0023 | 0.1198 | 0.9358 | True |
| Affinibrenneria | Lonsdalea | 0.322 | 0.2553 | -0.0902 | 0.7343 | False |
| Affinibrenneria | Musicola | 0.1782 | 0.9347 | -0.2769 | 0.6333 | False |
| Affinibrenneria | Pectobacterium | 0.511 | 0.0037 | 0.1035 | 0.9185 | True |
| Affinibrenneria | Samsonia | 0.1889 | 0.9748 | -0.3868 | 0.7645 | False |
| Brenneria | Dickeya | 0.5934 | 0 | 0.5171 | 0.6697 | True |
| Brenneria | Lonsdalea | 0.3877 | 0 | 0.2914 | 0.484 | True |
| Brenneria | Musicola | 0.2438 | 0.0142 | 0.0283 | 0.4593 | True |
| Brenneria | Pectobacterium | 0.5767 | 0 | 0.5031 | 0.6502 | True |
| Brenneria | Samsonia | 0.2545 | 0.5702 | -0.1586 | 0.6677 | False |
| Dickeya | Lonsdalea | -0.2057 | 0 | -0.2768 | -0.1347 | True |
| Dickeya | Musicola | -0.3496 | 0 | -0.5551 | -0.1441 | True |
| Dickeya | Pectobacterium | -0.0168 | 0.8188 | -0.0512 | 0.0177 | False |
| Dickeya | Samsonia | -0.3389 | 0.1866 | -0.7469 | 0.0691 | False |
| Lonsdalea | Musicola | -0.1439 | 0.4512 | -0.3576 | 0.0698 | False |
| Lonsdalea | Pectobacterium | 0.189 | 0 | 0.1209 | 0.257 | True |
| Lonsdalea | Samsonia | -0.1332 | 0.9769 | -0.5454 | 0.2791 | False |
| Musicola | Pectobacterium | 0.3328 | 0 | 0.1284 | 0.5373 | True |
| Musicola | Samsonia | 0.0107 | 1 | -0.4444 | 0.4658 | False |
| Pectobacterium | Samsonia | -0.3221 | 0.2413 | -0.7296 | 0.0854 | False |

### 4 Comparison of CAZome size

##### SI Figure 7: Boxplot overlaid by a scatter plot of the size of the CAZome

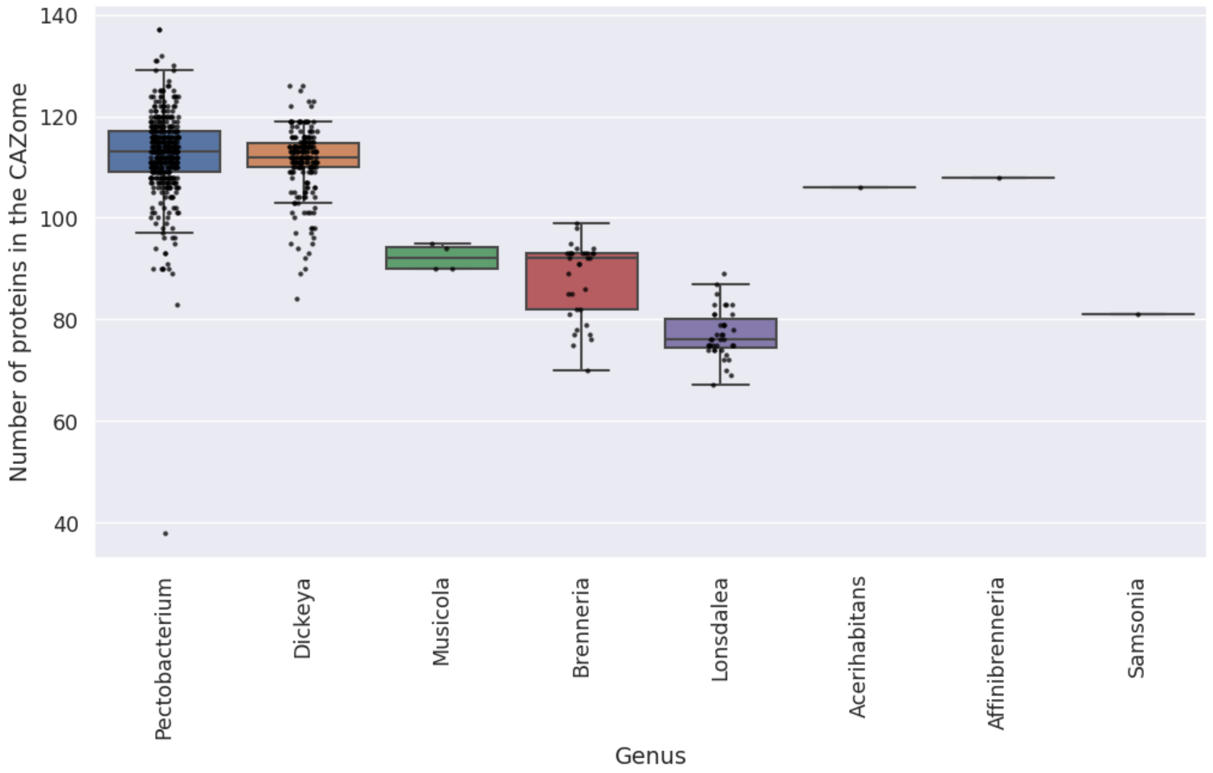


Figure 7: The number of CAZymes in the CAZome (i.e. the unique protein IDs in the CAZome) in each genome, grouped by genus with each point representing a unique genome, overlaying genomes are indicated with a darker shade.

#### SI Table 4: Tukey HDS post hoc test to identify statistically significant differences in mean CAZome size in soft and hard plant tissue targeting genomes

Output from a Tukey HDS post hoc test of the CAZome size (number of unique protein IDs in the CAZome) across soft plant tissue targeting genera (*Pectobacterium* and *Dickeya*) and hard plant tissue targeting genera (*Acerhabitans*, *Affinibrenneria*, *Brenneria*, *Lonsdalea*, and *Samsonia*). *Musicola* is labelled as 'unknown' owing to its preference of soft and hard plant tissues not having yet been established in the literature. A p-value (P-Adj, adjusted p-value) of 0.05 was used as a cut off rejecting the null hypothesis (NH). In addition the mean difference (MeanDiff) and lower and upper bounds of the confidence interval for the mean difference.

| Group1 | Group2 | MeanDiff | P-Adj | Lower | Upper | Reject NH |
|---|---|---|---|---|---|---|
| Hard | Soft | 29.4861 | 0 | 27.2598 | 31.7125 | True |
| Hard | Unknown | 9.57 | 0.0437 | 0.2108 | 18.9292 | True |
| Soft | Unknown | -19.91661 | 0 | -29.0639 | -10.7684 | True |

#### SI Table 5: Tukey HDS post hoc test to identify genera with statistically significantly different mean CAZome sizes

Output from a Tukey HDS post hoc test of the CAZome sizes (i.e. number of unique protein IDs) across the *Pectobacteriaceae* genera, performed using the Python StatsModels package (see Methods). A p-value (P-Adj, adjusted p-value) of 0.05 was used as a cut off rejecting the null hypothesis (NH). In addition the mean difference (MeanDiff) and lower and upper bounds of the confidence interval for the mean difference are indicated.

| Group1 | Group2 | MeanDiff | P-Adj | Lower | Upper | Reject NH |
|---|---|---|---|---|---|---|
| Acerihabitans | Affinibrenneria | 2 | 1 | -30.0935 | 34.0935 | False |
| Acerihabitans | Brenneria | -18.2121 | 0.2411 | -41.2469 | 4.8227 | False |
| Acerihabitans | Dickeya | 5.1602 | 0.9973 | -17.5883 | 27.9087 | False |
| Acerihabitans | Lonsdalea | -28.8462 | 0.0037 | -51.8288 | -5.8635 | True |
| Acerihabitans | Musicola | -13.75 | 0.7211 | -39.1221 | 11.6221 | False |
| Acerihabitans | Pectobacterium | 6.6458 | 0.987 | -16.0739 | 29.3656 | False |
| Acerihabitans | Samsonia | -25 | 0.2588 | -57.0935 | 7.0935 | False |
| Affinibrenneria | Brenneria | -20.2121 | 0.1344 | -43.2469 | 2.8227 | False |
| Affinibrenneria | Dickeya | 3.1602 | 0.9999 | -19.5883 | 25.9087 | False |
| Affinibrenneria | Lonsdalea | -30.8462 | 0.0013 | -53.8288 | -7.8635 | True |
| Affinibrenneria | Musicola | -15.75 | 0.5603 | -41.1221 | 9.6221 | False |
| Affinibrenneria | Pectobacterium | 4.6458 | 0.9986 | -18.0739 | 27.3656 | False |
| Affinibrenneria | Samsonia | -27 | 0.1735 | -59.0935 | 5.0935 | False |
| Brenneria | Dickeya | 23.3723 | 0 | 19.1172 | 27.6274 | True |
| Brenneria | Lonsdalea | -10.634 | 0 | -16.0016 | -5.2664 | True |
| Brenneria | Musicola | 4.4621 | 0.9504 | -7.5527 | 16.4769 | False |
| Brenneria | Pectobacterium | 24.858 | 0 | 20.7594 | 28.9565 | True |
| Brenneria | Samsonia | -6.7879 | 0.9864 | -29.8227 | 16.2469 | False |
| Dickeya | Lonsdalea | -34.0063 | 0 | -37.9693 | -30.0434 | True |
| Dickeya | Musicola | -18.9102 | 0 | -30.3666 | -7.4538 | True |
| Dickeya | Pectobacterium | 1.4856 | 0.2679 | -0.4358 | 3.4071 | False |
| Dickeya | Samsonia | -30.1602 | 0.0016 | -52.9087 | -7.4117 | True |
| Lonsdalea | Musicola | 15.0962 | 0.0032 | 3.1817 | 27.0106 | True |
| Lonsdalea | Pectobacterium | 35.492 | 0 | 31.6976 | 39.2863 | True |
| Lonsdalea | Samsonia | 3.8462 | 0.9996 | -19.1365 | 26.8288 | False |
| Musicola | Pectobacterium | 20.3958 | 0 | 8.9967 | 31.795 | True |
| Musicola | Samsonia | -11.25 | 0.88 | -36.6221 | 14.1221 | False |
| Pectobacterium | Samsonia | -31.6458 | 0.0007 | -54.3656 | -8.9261 | True |

### 5 Comparison of the number of CAZyme families

##### SI Figure 8: Boxplot overlaid by a scatter plot of the number of CAZy families in each genome

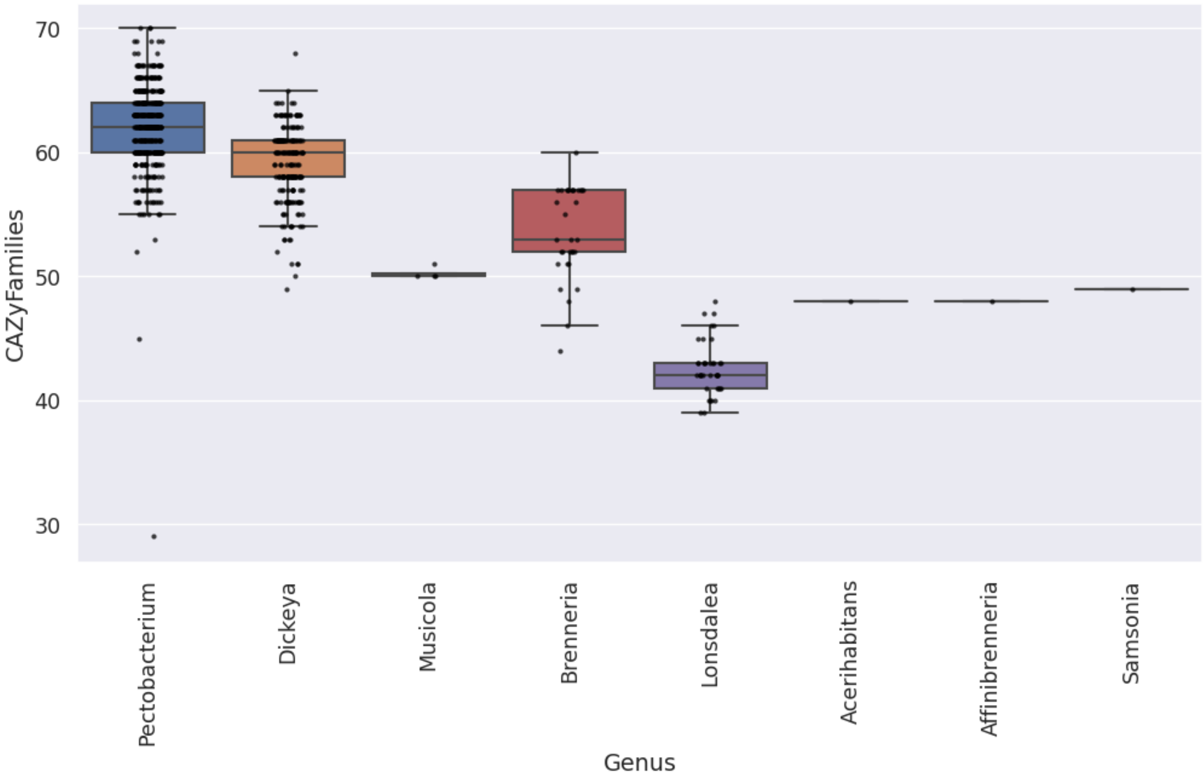


Figure 8: The number of CAZyme families in the CAZome of each genome, grouped by genus with each point representing a unique genome, overlaying genomes are indicated with a darker shade.

#### SI Table 6: Tukey HDS post hoc test to identify statistically significant differences in mean number of CAZy families in the CAZome in soft and hard plant tissue targeting genomes

Output from a Tukey HDS post hoc test of the mean number of CAZyme families in the CAZome across soft plant tissue targeting genera (*Pectobacterium* and *Dickeya*) and hard plant tissue targeting genera (*Acerhabitans*, *Affinibrenneria*, *Brenneria*, *Lonsdalea*, and *Samsonia*). *Musicola* is labelled as 'unknown' owing to its preference of soft and hard plant tissues not having yet been established in the literature. A p-value (P-Adj, adjusted p-value) of 0.05 was used as a cut off rejecting the null hypothesis (NH). In addition the mean difference (MeanDiff) and lower and upper bounds of the confidence interval for the mean difference.

| **Group1** | **Group2** | **MeanDiff** | **P-Adj** | **Lower** | **Upper** | **Reject NH** |
|---|---|---|---|---|---|---|
| Hard | Soft | 13.3936 | 0 | 12.593 | 14.528 | True |
| Hard | Unknown | 2.5433 | 0.4226 | -2.2253 | 7.312 | False |
| Soft | Unknown | -10.8503 | 0 | -15.5112 | -6.1894 | True |

#### SI Table 7: Tukey HDS post hoc test to identify genera with statistically significantly different mean number of CAZyme families

Output from a Tukey HDS post hoc test of the number of CAZy families in each CAZome, across the *Pectobacteriaceae* genera, performed using the Python StatsModels package (see Methods). A p-value (P-Adj, adjusted p-value) of 0.05 was used as a cut off rejecting the null hypothesis (NH). In addition the mean difference (MeanDiff) and lower and upper bounds of the confidence interval for the mean difference.

| Group1 | Group2 | MeanDiff | P-Adj | Lower | Upper | Reject NH |
|---|---|---|---|---|---|---|
| Acerihabitans | Affinibrenneria | 0 | 1 | -14.1792 | 14.1792 | False |
| Acerihabitans | Brenneria | 5.6667 | 0.6921 | -4.5103 | 15.8437 | False |
| Acerihabitans | Dickeya | 11.0485 | 0.0197 | 0.998 | 21.0991 | True |
| Acerihabitans | Lonsdalea | -5.3846 | 0.743 | -15.5386 | 4.7693 | False |
| Acerihabitans | Musicola | 2.25 | 0.9987 | -8.9597 | 13.4597 | False |
| Acerihabitans | Pectobacterium | 14.0787 | 0.0006 | 4.0409 | 24.1165 | True |
| Acerihabitans | Samsonia | 1 | 1 | -13.1792 | 15.1792 | False |
| Affinibrenneria | Brenneria | 5.6667 | 0.6921 | -4.5103 | 15.8437 | False |
| Affinibrenneria | Dickeya | 11.0485 | 0.0197 | 0.998 | 21.0991 | True |
| Affinibrenneria | Lonsdalea | -5.3846 | 0.743 | -15.5386 | 4.7693 | False |
| Affinibrenneria | Musicola | 2.25 | 0.9987 | -8.9597 | 13.4597 | False |
| Affinibrenneria | Pectobacterium | 14.0787 | 0.0006 | 4.0409 | 24.1165 | True |
| Affinibrenneria | Samsonia | 1 | 1 | -13.1792 | 15.1792 | False |
| Brenneria | Dickeya | 5.3819 | 0 | 3.5019 | 7.2618 | True |
| Brenneria | Lonsdalea | -11.0513 | 0 | -13.4227 | -8.6798 | True |
| Brenneria | Musicola | -3.4167 | 0.5122 | -8.7249 | 1.8916 | False |
| Brenneria | Pectobacterium | 8.412 | 0 | 6.6013 | 10.2228 | True |
| Brenneria | Samsonia | -4.6667 | 0.86 | -14.8437 | 5.5103 | False |
| Dickeya | Lonsdalea | -16.4332 | 0 | -18.184 | -14.6823 | True |
| Dickeya | Musicola | -8.7985 | 0 | -13.8601 | -3.737 | True |
| Dickeya | Pectobacterium | 3.0302 | 0 | 2.1812 | 3.8791 | True |
| Dickeya | Samsonia | -10.0485 | 0.0501 | -20.0991 | 0.002 | False |
| Lonsdalea | Musicola | 7.6346 | 0.0003 | 2.3707 | 12.8985 | True |
| Lonsdalea | Pectobacterium | 19.4633 | 0 | 17.7869 | 21.1397 | True |
| Lonsdalea | Samsonia | 6.3846 | 0.5434 | -3.7693 | 16.5386 | False |
| Musicola | Pectobacterium | 11.8287 | 0 | 6.7924 | 16.865 | True |
| Musicola | Samsonia | -1.25 | 1 | -12.4597 | 9.9597 | False |
| Pectobacterium | Samsonia | -13.0787 | 0.0021 | -23.1165 | -3.0409 | True |

### 6 Comparison of CAZy class frequencies

#### SI Table 8: Two-way ANOVA of CAZy class frequencies

Output from a two-way ANOVA (performed using the Python StatsModels package, see Methods) testing for statistically significantly differences in the mean frequencies between the CAZy classes and *Pectobacteriaceae* genera. A PR($\geq$F) value (the p-value) of 0.05 was used as the cut-off.

| Category | Sum of squares | Degrees of freedom | F value | PR(>F) |
|---|---|---|---|---|
| **Genus** | 1.37E4 | 7 | 137.43 | 4.38E-183 |
| **CAZy Class** | 1.11E6 | 5 | 15606.45 | 0 |
| **Genus:CAZy Class** | 2.4E4 | 35 | 48.05 | 1.70E-276 |
| **Residuals** | 6.07e+04 | 4254 | NA | NA |

#### SI Table 9: Tukey HSD test of Glycoside Hydrolase (GH) CAZy class frequencies

Output from a Tukey HDS post hoc test of the number of CAZymes in the Glycoside Hydrolase (GH) CAZy class across the *Pectobacteriaceae* genera, performed using the Python StatsModels package (see Methods). A p-value (P-Adj, adjusted p-value) of 0.05 was used as a cut off rejecting the null hypothesis (NH). In addition the mean difference (MeanDiff) and lower and upper bounds of the confidence interval for the mean difference.

| Group 1 | Group 2 | MeanDiff | Adjusted P-value | Lower | Upper | Reject NH |
|---|---|---|---|---|---|---|
| Acerihabitans | Affinibrenneria | 8 | 0.9884 | -19.8655 | 35.8655 | False |
| Acerihabitans | Brenneria | -8.5152 | 0.901 | -28.5153 | 11.485 | False |
| Acerihabitans | Dickeya | -7.6699 | 0.9374 | -27.4215 | 12.0817 | False |
| Acerihabitans | Lonsdalea | -20.1538 | 0.0458 | -40.1087 | -0.199 | True |
| Acerihabitans | Musicola | -14 | 0.5293 | -36.0296 | 8.0296 | False |
| Acerihabitans | Pectobacterium | -0.1551 | 1 | -19.8818 | 19.5716 | False |
| Acerihabitans | Samsonia | -17 | 0.5827 | -44.8655 | 10.8655 | False |
| Affinibrenneria | Brenneria | -16.5152 | 0.1927 | -36.5153 | 3.485 | False |
| Affinibrenneria | Dickeya | -15.6699 | 0.2371 | -35.4215 | 4.0817 | False |
| Affinibrenneria | Lonsdalea | -28.1538 | 0.0005 | -48.1087 | -8.199 | True |
| Affinibrenneria | Musicola | -22 | 0.0506 | -44.0296 | 0.0296 | False |
| Affinibrenneria | Pectobacterium | -8.1551 | 0.9142 | -27.8818 | 11.5716 | False |
| Affinibrenneria | Samsonia | -25 | 0.1161 | -52.8655 | 2.8655 | False |
| Brenneria | Dickeya | 0.8452 | 0.9971 | -2.8493 | 4.5398 | False |
| Brenneria | Lonsdalea | -11.6387 | 0 | -16.2992 | -6.9782 | True |
| Brenneria | Musicola | -5.4848 | 0.7514 | -15.9168 | 4.9471 | False |
| Brenneria | Pectobacterium | 8.3601 | 0 | 4.8015 | 11.9187 | True |
| Brenneria | Samsonia | -8.4848 | 0.9027 | -28.485 | 11.5153 | False |
| Dickeya | Lonsdalea | -12.4839 | 0 | -15.9248 | -9.0431 | True |
| Dickeya | Musicola | -6.3301 | 0.5274 | -16.2772 | 3.617 | False |
| Dickeya | Pectobacterium | 7.5148 | 0 | 5.8465 | 9.1832 | True |
| Dickeya | Samsonia | -9.3301 | 0.8403 | -29.0817 | 10.4215 | False |
| Lonsdalea | Musicola | 6.1538 | 0.6146 | -4.191 | 16.4987 | False |
| Lonsdalea | Pectobacterium | 19.9988 | 0 | 16.7043 | 23.2932 | True |
| Lonsdalea | Samsonia | 3.1538 | 0.9997 | -16.801 | 23.1087 | False |
| Musicola | Pectobacterium | 13.8449 | 0.0006 | 3.9475 | 23.7423 | True |
| Musicola | Samsonia | -3 | 0.9999 | -25.0296 | 19.0296 | False |
| Pectobacterium | Samsonia | -16.8449 | 0.159 | -36.5716 | 2.8818 | False |

#### SI Table 10: Tukey HSD test of Glycosyltransferase (GT) CAZy class frequencies

Output from a Tukey HDS post hoc test of the number of CAZymes in the Glycosyltransferase (GT) CAZy class across the *Pectobacteriaceae* genera, performed using the Python StatsModels package (see Methods). A p-value (P-Adj, adjusted p-value) of 0.05 was used as a cut off rejecting the null hypothesis (NH). In addition the mean difference (MeanDiff) and lower and upper bounds of the confidence interval for the mean difference.

| Group 1 | Group 2 | MeanDiff | Adjusted P-value | Lower | Upper | Reject NH |
|---|---|---|---|---|---|---|
| Acerihabitans | Affinibrenneria | -9 | 0.9555 | -33.7332 | 15.7332 | False |
| Acerihabitans | Brenneria | -11.6667 | 0.484 | -29.4187 | 6.0853 | False |
| Acerihabitans | Dickeya | -6.0437 | 0.9668 | -23.5751 | 11.4877 | False |
| Acerihabitans | Lonsdalea | -11.5385 | 0.4959 | -29.2502 | 6.1733 | False |
| Acerihabitans | Musicola | -13 | 0.4682 | -32.5533 | 6.5533 | False |
| Acerihabitans | Pectobacterium | -11.7361 | 0.4571 | -29.2453 | 5.7731 | False |
| Acerihabitans | Samsonia | -19 | 0.2758 | -43.7332 | 5.7332 | False |
| Affinibrenneria | Brenneria | -2.6667 | 0.9998 | -20.4187 | 15.0853 | False |
| Affinibrenneria | Dickeya | 2.9563 | 0.9996 | -14.5751 | 20.4877 | False |
| Affinibrenneria | Lonsdalea | -2.5385 | 0.9999 | -20.2502 | 15.1733 | False |
| Affinibrenneria | Musicola | -4 | 0.9986 | -23.5533 | 15.5533 | False |
| Affinibrenneria | Pectobacterium | -2.7361 | 0.9998 | -20.2453 | 14.7731 | False |
| Affinibrenneria | Samsonia | -10 | 0.9232 | -34.7332 | 14.7332 | False |
| Brenneria | Dickeya | 5.623 | 0 | 2.3437 | 8.9022 | True |
| Brenneria | Lonsdalea | 0.1282 | 1 | -4.0084 | 4.2648 | False |
| Brenneria | Musicola | -1.3333 | 0.9999 | -10.5926 | 7.926 | False |
| Brenneria | Pectobacterium | -0.0694 | 1 | -3.228 | 3.0891 | False |
| Brenneria | Samsonia | -7.3333 | 0.9145 | -25.0853 | 10.4187 | False |
| Dickeya | Lonsdalea | -5.4948 | 0 | -8.5489 | -2.4407 | True |
| Dickeya | Musicola | -6.9563 | 0.2452 | -15.7853 | 1.8727 | False |
| Dickeya | Pectobacterium | -5.6924 | 0 | -7.1732 | -4.2116 | True |
| Dickeya | Samsonia | -12.9563 | 0.3252 | -30.4877 | 4.5751 | False |
| Lonsdalea | Musicola | -1.4615 | 0.9997 | -10.6435 | 7.7204 | False |
| Lonsdalea | Pectobacterium | -0.1976 | 1 | -3.1218 | 2.7265 | False |
| Lonsdalea | Samsonia | -7.4615 | 0.9059 | -25.1733 | 10.2502 | False |
| Musicola | Pectobacterium | 1.2639 | 0.9999 | -7.521 | 10.0488 | False |
| Musicola | Samsonia | -6 | 0.9828 | -25.5533 | 13.5533 | False |
| Pectobacterium | Samsonia | -7.2639 | 0.9127 | -24.7731 | 10.2453 | False |

#### SI Table 11: Tukey HSD test of Carbohydrate Esterase (CE) CAZy class frequencies

Output from a Tukey HDS post hoc test of the number of CAZymes in the Carbohydrate Esterase (CE) CAZy class across the *Pectobacteriaceae* genera, performed using the Python StatsModels package (see Methods). A p-value (P-Adj, adjusted p-value) of 0.05 was used as a cut off rejecting the null hypothesis (NH). In addition the mean difference (MeanDiff) and lower and upper bounds of the confidence interval for the mean difference.

| **Group 1** | **Group 2** | **MeanDiff** | **P-Adj** | **Lower** | **Upper** | **Reject NH** |
|---|---|---|---|---|---|---|
| Acerihabitans | Affinibrenneria | 2 | 0.8959 | -2.6483 | 6.6483 | False |
| Acerihabitans | Brenneria | -0.697 | 0.9984 | -4.0332 | 2.6393 | False |
| Acerihabitans | Dickeya | 2.2767 | 0.4155 | -1.0181 | 5.5715 | False |
| Acerihabitans | Lonsdalea | -1.8462 | 0.6964 | -5.1749 | 1.4826 | False |
| Acerihabitans | Musicola | 1 | 0.9916 | -2.6748 | 4.6748 | False |
| Acerihabitans | Pectobacterium | 2.2176 | 0.4497 | -1.0731 | 5.5082 | False |
| Acerihabitans | Samsonia | 3 | 0.5085 | -1.6483 | 7.6483 | False |
| Affinibrenneria | Brenneria | -2.697 | 0.2157 | -6.0332 | 0.6393 | False |
| Affinibrenneria | Dickeya | 0.2767 | 1 | -3.0181 | 3.5715 | False |
| Affinibrenneria | Lonsdalea | -3.8462 | 0.0111 | -7.1749 | -0.5174 | True |
| Affinibrenneria | Musicola | -1 | 0.9916 | -4.6748 | 2.6748 | False |
| Affinibrenneria | Pectobacterium | 0.2176 | 1 | -3.0731 | 3.5082 | False |
| Affinibrenneria | Samsonia | 1 | 0.998 | -3.6483 | 5.6483 | False |
| Brenneria | Dickeya | 2.9737 | 0 | 2.3574 | 3.59 | True |
| Brenneria | Lonsdalea | -1.1492 | 0.0002 | -1.9266 | -0.3718 | True |
| Brenneria | Musicola | 1.697 | 0.062 | -0.0432 | 3.4371 | False |
| Brenneria | Pectobacterium | 2.9146 | 0 | 2.3209 | 3.5082 | True |
| Brenneria | Samsonia | 3.697 | 0.018 | 0.3607 | 7.0332 | True |
| Dickeya | Lonsdalea | -4.1229 | 0 | -4.6968 | -3.5489 | True |
| Dickeya | Musicola | -1.2767 | 0.2739 | -2.936 | 0.3826 | False |
| Dickeya | Pectobacterium | -0.0591 | 0.9982 | -0.3374 | 0.2192 | False |
| Dickeya | Samsonia | 0.7233 | 0.9978 | -2.5715 | 4.0181 | False |
| Lonsdalea | Musicola | 2.8462 | 0 | 1.1205 | 4.5718 | True |
| Lonsdalea | Pectobacterium | 4.0637 | 0 | 3.5142 | 4.6133 | True |
| Lonsdalea | Samsonia | 4.8462 | 0.0003 | 1.5174 | 8.1749 | True |
| Musicola | Pectobacterium | 1.2176 | 0.3279 | -0.4334 | 2.8686 | False |
| Musicola | Samsonia | 2 | 0.7166 | -1.6748 | 5.6748 | False |
| Pectobacterium | Samsonia | 0.7824 | 0.9963 | -2.5082 | 4.0731 | False |

#### SI Table 12: Tukey HSD test of Carbohydrate Binding Module (CBM) CAZy class frequencies

Output from a Tukey HDS post hoc test of the number of CAZymes in the Carbohydrate Binding Module (CBM) CAZy class across the *Pectobacteriaceae* genera, performed using the Python StatsModels package (see Methods). A p-value (P-Adj, adjusted p-value) of 0.05 was used as a cut off rejecting the null hypothesis (NH). In addition the mean difference (MeanDiff) and lower and upper bounds of the confidence interval for the mean difference.

| Group 1 | Group 2 | MeanDiff | P-Adj | Lower | Upper | Reject NH |
|---|---|---|---|---|---|---|
| Acerihabitans | Affinibrenneria | -1 | 0.9999 | -8.6335 | 6.6335 | False |
| Acerihabitans | Brenneria | -1.6364 | 0.9853 | -7.1152 | 3.8425 | False |
| Acerihabitans | Dickeya | 1.4272 | 0.993 | -3.9836 | 6.838 | False |
| Acerihabitans | Lonsdalea | -2.2564 | 0.9148 | -7.7229 | 3.2101 | False |
| Acerihabitans | Musicola | 0 | 1 | -6.0348 | 6.0348 | False |
| Acerihabitans | Pectobacterium | 3.1204 | 0.6506 | -2.2836 | 8.5243 | False |
| Acerihabitans | Samsonia | -1 | 0.9999 | -8.6335 | 6.6335 | False |
| Affinibrenneria | Brenneria | -0.6364 | 1 | -6.1152 | 4.8425 | False |
| Affinibrenneria | Dickeya | 2.4272 | 0.8734 | -2.9836 | 7.838 | False |
| Affinibrenneria | Lonsdalea | -1.2564 | 0.997 | -6.7229 | 4.2101 | False |
| Affinibrenneria | Musicola | 1 | 0.9996 | -5.0348 | 7.0348 | False |
| Affinibrenneria | Pectobacterium | 4.1204 | 0.2852 | -1.2836 | 9.5243 | False |
| Affinibrenneria | Samsonia | 0 | 1 | -7.6335 | 7.6335 | False |
| Brenneria | Dickeya | 3.0635 | 0 | 2.0515 | 4.0756 | True |
| Brenneria | Lonsdalea | -0.62 | 0.8199 | -1.8967 | 0.6566 | False |
| Brenneria | Musicola | 1.6364 | 0.6604 | -1.2214 | 4.4941 | False |
| Brenneria | Pectobacterium | 4.7567 | 0 | 3.7819 | 5.7316 | True |
| Brenneria | Samsonia | 0.6364 | 1 | -4.8425 | 6.1152 | False |
| Dickeya | Lonsdalea | -3.6836 | 0 | -4.6262 | -2.741 | True |
| Dickeya | Musicola | -1.4272 | 0.755 | -4.1521 | 1.2977 | False |
| Dickeya | Pectobacterium | 1.6932 | 0 | 1.2362 | 2.1502 | True |
| Dickeya | Samsonia | -2.4272 | 0.8734 | -7.838 | 2.9836 | False |
| Lonsdalea | Musicola | 2.2564 | 0.2329 | -0.5775 | 5.0903 | False |
| Lonsdalea | Pectobacterium | 5.3768 | 0 | 4.4743 | 6.2793 | True |
| Lonsdalea | Samsonia | 1.2564 | 0.997 | -4.2101 | 6.7229 | False |
| Musicola | Pectobacterium | 3.1204 | 0.0116 | 0.4091 | 5.8317 | True |
| Musicola | Samsonia | -1 | 0.9996 | -7.0348 | 5.0348 | False |
| Pectobacterium | Samsonia | -4.1204 | 0.2852 | -9.5243 | 1.2836 | False |

#### SI Table 13: Tukey HSD test of Carbohydrate Binding Module (CBM) CAZy class frequencies between soft and hard plant tissue targeting genera

Output from a Tukey HDS post hoc test of the number of CAZymes in the Carbohydrate Binding Module (CBM) CAZy class across soft and hard plant tissue targeting genera in *Pectobacteriaceae* and *Musicola*, performed using the Python StatsModels package (see Methods). A p-value (P-Adj, adjusted p-value) of 0.05 was used as a cut off rejecting the null hypothesis (NH). In addition the mean difference (MeanDiff) and lower and upper bounds of the confidence interval for the mean difference are shown.

| **Group 1** | **Group 2** | **MeanDiff** | **P-Adj** | **Lower** | **Upper** | **Reject NH** |
|---|---|---|---|---|---|---|
| Hard | Soft | 4.4937 | 0 | 3.9416 | 5.0457 | True |
| Hard | Unknown | 1.92 | 0.1275 | -0.4008 | 4.2408 | False |
| Soft | Unknown | -2.5737 | 0.0215 | -4.842 | -0.3053 | True |

#### SI Table 14: Tukey HSD test of Auxiliary Activity (AA) CAZy class frequencies

Output from a Tukey HDS post hoc test of the number of CAZymes in the Auxiliary Activity (AA) CAZy class across the *Pectobacteriaceae* genera, performed using the Python StatsModels package (see Methods). A p-value (P-Adj, adjusted p-value) of 0.05 was used as a cut off rejecting the null hypothesis (NH). In addition the mean difference (MeanDiff) and lower and upper bounds of the confidence interval for the mean difference are shown.

| **Group 1** | **Group 2** | **MeanDiff** | **P-Adj** | **Lower** | **Upper** | **Reject NH** |
|---|---|---|---|---|---|---|
| Acerihabitans | Affinibrenneria | 0 | 1 | -1.7994 | 1.7994 | False |
| Acerihabitans | Brenneria | 0.2424 | 0.9992 | -1.0491 | 1.5339 | False |
| Acerihabitans | Dickeya | 0.4126 | 0.9767 | -0.8628 | 1.6881 | False |
| Acerihabitans | Lonsdalea | 0 | 1 | -1.2886 | 1.2886 | False |
| Acerihabitans | Musicola | 0 | 1 | -1.4225 | 1.4225 | False |
| Acerihabitans | Pectobacterium | 0.8843 | 0.4092 | -0.3896 | 2.1581 | False |
| Acerihabitans | Samsonia | 1 | 0.6942 | -0.7994 | 2.7994 | False |
| Affinibrenneria | Brenneria | 0.2424 | 0.9992 | -1.0491 | 1.5339 | False |
| Affinibrenneria | Dickeya | 0.4126 | 0.9767 | -0.8628 | 1.6881 | False |
| Affinibrenneria | Lonsdalea | 0 | 1 | -1.2886 | 1.2886 | False |
| Affinibrenneria | Musicola | 0 | 1 | -1.4225 | 1.4225 | False |
| Affinibrenneria | Pectobacterium | 0.8843 | 0.4092 | -0.3896 | 2.1581 | False |
| Affinibrenneria | Samsonia | 1 | 0.6942 | -0.7994 | 2.7994 | False |
| Brenneria | Dickeya | 0.1702 | 0.3721 | -0.0684 | 0.4088 | False |
| Brenneria | Lonsdalea | -0.2424 | 0.2197 | -0.5434 | 0.0585 | False |
| Brenneria | Musicola | -0.2424 | 0.958 | -0.9161 | 0.4312 | False |
| Brenneria | Pectobacterium | 0.6418 | 0 | 0.412 | 0.8716 | True |
| Brenneria | Samsonia | 0.7576 | 0.6317 | -0.5339 | 2.0491 | False |
| Dickeya | Lonsdalea | -0.4126 | 0 | -0.6348 | -0.1904 | True |
| Dickeya | Musicola | -0.4126 | 0.5148 | -1.055 | 0.2297 | False |
| Dickeya | Pectobacterium | 0.4716 | 0 | 0.3639 | 0.5794 | True |
| Dickeya | Samsonia | 0.5874 | 0.8573 | -0.6881 | 1.8628 | False |
| Lonsdalea | Musicola | 0 | 1 | -0.668 | 0.668 | False |
| Lonsdalea | Pectobacterium | 0.8843 | 0 | 0.6715 | 1.097 | True |
| Lonsdalea | Samsonia | 1 | 0.2634 | -0.2886 | 2.2886 | False |
| Musicola | Pectobacterium | 0.8843 | 0.0008 | 0.2451 | 1.5234 | True |
| Musicola | Samsonia | 1 | 0.392 | -0.4225 | 2.4225 | False |
| Pectobacterium | Samsonia | 0.1157 | 1 | -1.1581 | 1.3896 | False |

#### SI Table 15: Tukey HSD test of Auxiliary Activity (AA) CAZy class frequencies between soft and hard plant tissue targeting genera

Output from a Tukey HDS post hoc test of the number of CAZymes in the Auxiliary Activity (AA) CAZy class across soft and hard plant tissue targeting genera in *Pectobacteriaceae* and *Musicola*, performed using the Python StatsModels package (see Methods). A p-value (P-Adj, adjusted p-value) of 0.05 was used as a cut off rejecting the null hypothesis (NH). In addition the mean difference (MeanDiff) and lower and upper bounds of the confidence interval for the mean difference are shown.

| Group 1 | Group 2 | MeanDiff | P-Adj | Lower | Upper | Reject NH |
|---|---|---|---|---|---|---|
| Hard | Soft | 0.612 | 0 | 0.4775 | 0.7464 | True |
| Hard | Unknown | -0.12 | 0.872 | -0.6853 | 0.4453 | False |
| Soft | Unknown | -0.732 | 0.0055 | -1.2845 | -0.1794 | True |

#### SI Table 16: Tukey HSD test of Polysaccharide Lyase (PL) CAZy class frequencies between soft and hard plant tissue targeting genera

Output from a Tukey HDS post hoc test of the number of CAZymes in the Polysaccharide Lyase (PL) CAZy class across soft and hard plant tissue targeting genera in *Pectobacteriaceae* and *Musicola*, performed using the Python StatsModels package (see Methods). A p-value (P-Adj, adjusted p-value) of 0.05 was used as a cut off rejecting the null hypothesis (NH). In addition the mean difference (MeanDiff) and lower and upper bounds of the confidence interval for the mean difference are shown.

| Group 1 | Group 2 | MeanDiff | P-Adj | Lower | Upper | Reject NH |
|---|---|---|---|---|---|---|
| Hard | Soft | 11.5549 | 0 | 10.8223 | 12.2874 | True |
| Hard | Unknown | 7.2767 | 0 | 4.1971 | 10.3562 | True |
| Soft | Unknown | -4.2782 | 0.0026 | -7.2882 | -1.2683 | True |

#### SI Table 17: Tukey HSD test of Polysaccharide Lyase (PL) CAZy class frequencies

Output from a Tukey HDS post hoc test of the number of CAZymes in the Polysaccharide Lyase (PL) CAZy class across the *Pectobacteriaceae* genera, performed using the Python StatsModels package (see Methods). A p-value (P-Adj, adjusted p-value) of 0.05 was used as a cut off rejecting the null hypothesis (NH). In addition the mean difference (MeanDiff) and lower and upper bounds of the confidence interval for the mean difference are shown.

| Group 1 | Group 2 | MeanDiff | P-Adj | Lower | Upper | Reject NH |
|---|---|---|---|---|---|---|
| Acerihabitans | Affinibrenneria | 0 | 1 | -10.5548 | 10.5548 | False |
| Acerihabitans | Brenneria | 3.2424 | 0.8984 | -4.3332 | 10.8181 | False |
| Acerihabitans | Dickeya | 15.6019 | 0 | 8.1204 | 23.0834 | True |
| Acerihabitans | Lonsdalea | 2.7949 | 0.9515 | -4.7636 | 10.3534 | False |
| Acerihabitans | Musicola | 10.25 | 0.005 | 1.9057 | 18.5943 | True |
| Acerihabitans | Pectobacterium | 14.0162 | 0 | 6.5442 | 21.4882 | True |
| Acerihabitans | Samsonia | 7 | 0.4716 | -3.5548 | 17.5548 | False |
| Affinibrenneria | Brenneria | 3.2424 | 0.8984 | -4.3332 | 10.8181 | False |
| Affinibrenneria | Dickeya | 15.6019 | 0 | 8.1204 | 23.0834 | True |
| Affinibrenneria | Lonsdalea | 2.7949 | 0.9515 | -4.7636 | 10.3534 | False |
| Affinibrenneria | Musicola | 10.25 | 0.005 | 1.9057 | 18.5943 | True |
| Affinibrenneria | Pectobacterium | 14.0162 | 0 | 6.5442 | 21.4882 | True |
| Affinibrenneria | Samsonia | 7 | 0.4716 | -3.5548 | 17.5548 | False |
| Brenneria | Dickeya | 12.3595 | 0 | 10.9601 | 13.7589 | True |
| Brenneria | Lonsdalea | -0.4476 | 0.9945 | -2.2128 | 1.3177 | False |
| Brenneria | Musicola | 7.0076 | 0 | 3.0562 | 10.959 | True |
| Brenneria | Pectobacterium | 10.7738 | 0 | 9.4259 | 12.1217 | True |
| Brenneria | Samsonia | 3.7576 | 0.8033 | -3.8181 | 11.3332 | False |
| Dickeya | Lonsdalea | -12.8071 | 0 | -14.1104 | -11.5037 | True |
| Dickeya | Musicola | -5.3519 | 0.0005 | -9.1197 | -1.5842 | True |
| Dickeya | Pectobacterium | -1.5857 | 0 | -2.2177 | -0.9538 | True |
| Dickeya | Samsonia | -8.6019 | 0.0118 | -16.0834 | -1.1204 | True |
| Lonsdalea | Musicola | 7.4551 | 0 | 3.5367 | 11.3735 | True |
| Lonsdalea | Pectobacterium | 11.2213 | 0 | 9.9735 | 12.4692 | True |
| Lonsdalea | Samsonia | 4.2051 | 0.693 | -3.3534 | 11.7636 | False |
| Musicola | Pectobacterium | 3.7662 | 0.048 | 0.0173 | 7.5151 | True |
| Musicola | Samsonia | -3.25 | 0.9364 | -11.5943 | 5.0943 | False |
| Pectobacterium | Samsonia | -7.0162 | 0.0837 | -14.4882 | 0.4558 | False |

### 7 Principal Component Analysis (PCA) of CAZy family frequencies

#### SI Figure 9: Variance captured by principal component analysis of *Pectobacteriaceae* CAZy family frequencies

The number of unique protein accessions was calculated for each CAZy family, per genome. Dimensional reduction was performed on these CAZy family frequencies. This figure plots the cumulative frequency across the computed principal components (PCs). The scree plot of this figure plots the proportion of variance in the original data set that is captured by each PC. The *Pectobacteriaceae* genomes were projected onto principal components (PCs) PC1 and PC2, and *Dickeya* and *Pectobacterium* genomes located in the positive X-axis are annotated in the figure.

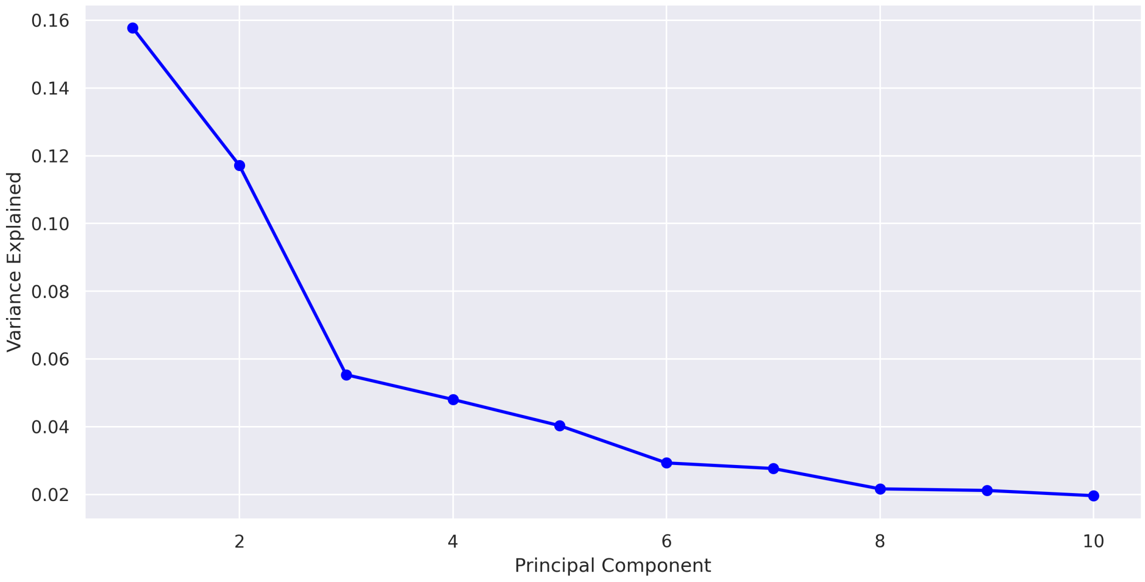


(a) Scree plot

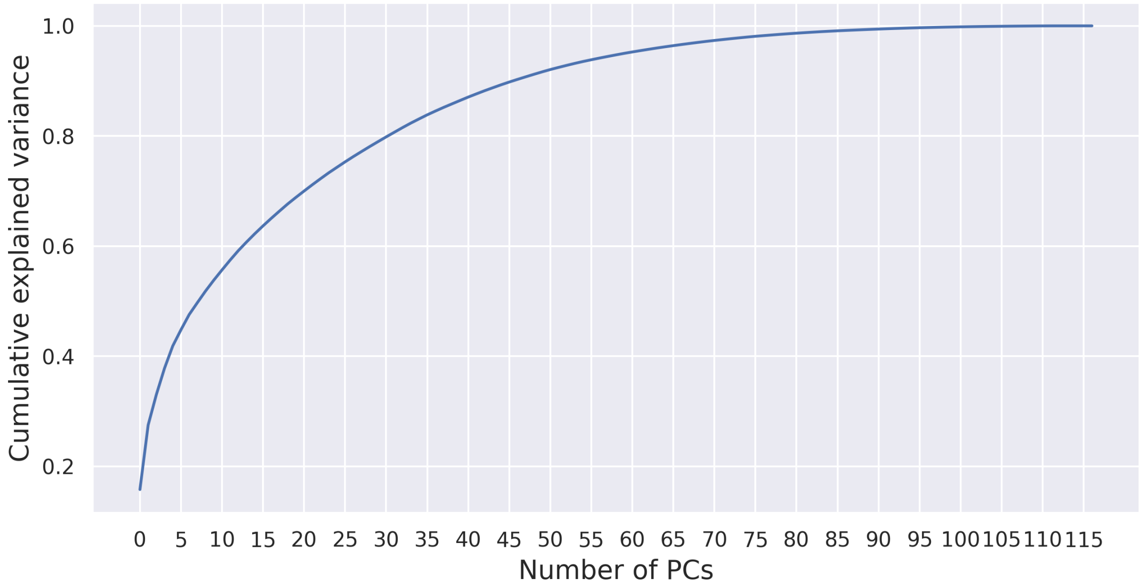


(b) Cumulative explained variance

Figure 9: Principal component analysis (PCA) of the CAZy family frequencies in *Pectobacteriaceae* genomes, plotting [A] the cumulative frequency across all computed principal components (PCs). [B] Scree plot plotting the fraction of variance captured by each PC.

#### SI Figure 10: Principal component analysis of *Pectobacteriaceae* CAZy family frequencies PC1 and PC2

The number of unique protein accessions was calculated for each CAZy family, per genome. Dimensional reduction was performed on these CAZy family frequencies. The *Pectobacteriaceae* genomes were projected onto principal components (PCs) PC1 and PC2, annotating *Dickeya* genomes that cluster with *Musicola* genomes (figure 10).

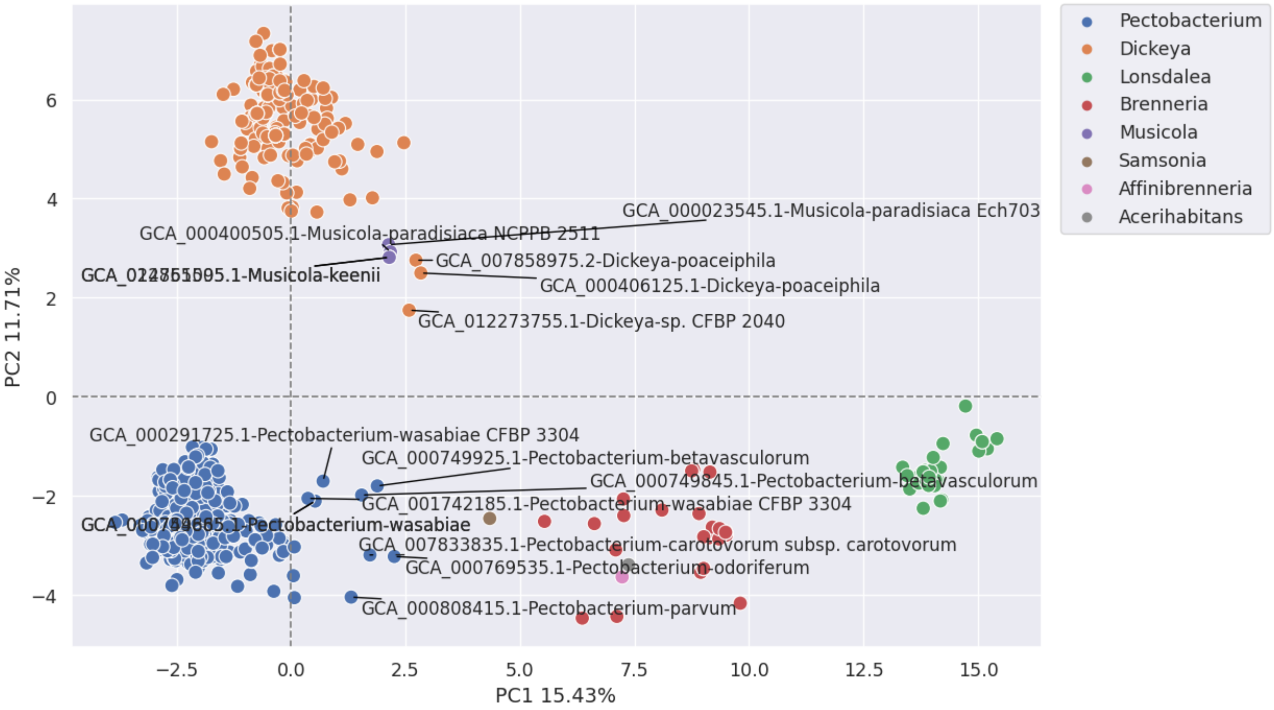


Figure 10: Principal component analysis (PCA) of the CAZy family frequencies in *Pectobacteriaceae* genomes, projecting genomes onto principal components (PCs) PC1 and PC2, colour coding genomes by genus classification. *Dickeya* genomes that cluster with *Musicola* genomes are annotated, as well as *Pectobacterium* genomes that are placed in the positive PC1-axis.

#### SI Table 18: Phenotypic comparison of *Musicola* and *Dickeya poaceiphila*

Carbon sources highlighted in green indicate conditions where both *D. poaceiphila* display a different phenotype to both *Musicola* genomes. Carbon sources/conditions highlighted in yellow indicate conditions where one of genomes shows a different phenotype to the other three.

| Carbon source | Strains | 3937 | A6065T (CFBP 4178T ) | A3967T (CFBP 8732T ) | NCPPB 569T | CFBP 2040 | 3937 |
|---|---|---|---|---|---|---|---|
| Both genomes differs from the other 4 | Species | *D. dadantii* | *M. paradisiaca* | *M. keenii* | *D.poaceiphila* | *D.poaceiphila* | *D.dadantii* |
| At least one genome differs from *D.popoaceiphila* | Reference | Hugouvieux-Cotte-Pattat et al., 2021; https://doi.org/10.1099/ijsem.0.005037 | | | Hugouvieux-Cotte-Pattat et al., 2020; https://doi.org/10.1099/ijsem.0.004306 | | |
| L-Arabinose | | + | - | + | + | + | + |
| N-Acetyl-D-glucosamine | | + | + | + | + | + | + |
| D-Saccharic acid (glucaric acid) | | + | - | + | + | + | + |
| Succinic acid | | w | w | + | + | + | w |
| D-Galactose | | + | - | + | w | + | + |
| L-Aspartic acid | | + | w | + | + | + | + |
| L-Proline | | - | - | - | - | - | - |
| D-Alanine | | - | - | - | - | - | - |
| D-Trehalose | | - | - | - | - | - | - |
| D-Mannose | | + | + | + | + | + | + |
| Dulcitol | | - | - | - | - | - | - |
| D-Serine | | - | - | - | - | - | - |
| D-Sorbitol | | - | - | - | - | - | - |
| Glycerol | | + | w | + | w | + | + |
| L-Fucose | | - | - | - | - | - | - |
| D-Glucuronic acid | | - | w | + | - | w | - |
| D-Gluconic acid | | + | + | + | - | - | + |
| D,L-α-Glycerol-phosphate | | w | w | w | w | w | w |
| D-Xylose | | + | - | + | + | + | + |
| L-Lactic acid | | w | - | - | + | + | w |
| Formic acid | | w | - | - | + | + | w |
| D-Mannitol | | + | - | - | + | + | + |
| L-Glutamic acid | | w | - | - | w | w | w |
| D-Glucose-6-phosphate | | + | - | + | - | - | + |
| D-Galactonic acid-γ-lactone | | - | - | - | - | - | - |
| D,L-Malic acid | | + | w | + | + | + | + |
| D-Ribose | | + | - | + | + | + | w |
| Tween 20 | | - | - | - | - | - | - |
| L-Rhamnose | | - | - | - | + | + | - |
| D-Fructose | | + | + | + | + | + | + |
| Acetic acid | | w | - | - | w | w | w |
| α-D-Glucose | | + | + | + | + | + | + |
| Maltose | | - | - | - | - | - | - |
| D-Melibiose | | + | + | - | + | + | + |
| Thymidine | | - | - | - | w | w | - |
| L-Asparagine | | + | w | + | + | + | + |
| D-Aspartic acid | | + | - | - | w | + | + |
| D-Glucosaminic acid | | - | - | - | - | - | - |
| 1,2-Propanediol | | - | - | - | - | - | - |
| Tween 40 | | - | - | - | - | - | - |
| α-Keto-glutaric acid | | - | - | - | - | - | - |
| α-Keto-butyric acid | | - | - | - | - | - | - |
| α-Methyl-D-galactoside | | + | - | - | w | w | + |
| α-D-Lactose | | - | - | - | w | + | - |
| Lactulose | | - | - | - | - | - | - |

#### SI Table 19: Genomes with potentially less rich CAZomes

Table listing the assembly status and CheckM analysis of the genomic sequence completeness as reported in the NCBI Assembly database (October 2023) for genomes that contained fewer families than genomes from the same genus (as identified in SI figures 10-13).

Assembly status and CheckM genomic assembly analysis reported in the NCBI Assembly database (October 2023)

| Assembly Accession | Genus | Species | Assembly status | CheckM analysis (% (percentile) |
|---|---|---|---|---|
| GCA_029023745.1 | *Pectobacterium* | *colocasium* PL155 | Complete | NA |
| GCA_021907015.1 | *Pectobacterium* | sp. PL152 | Complete | NA |
| GCA_000808415.1 | *Pectobacterium* | *parvum* Y1 | Scaffold | 81.03 (13th) |
| GCA_004137815.1 | *Pectobacterium* | *zantedeschiae* 2M | Scaffold | 89.31 (33rd) |
| GCA_012273755.1 | *Dickeya* | sp. CFBP 2040 | Contig | 90.95 (100th) |
| GCA_000406125.1 | *Dickeya* | *poaceiphila* NCPPB 569 | Chromosome | 89.80 (50th) |
| GCA_013168485.1 | *Dickeya* | *dadantii* Aka1-1 | Contig | 83.55 (11th) |
| GCA_007858975.2 | *Dickeya* | *poaceiphila* NCPPB 569 | Complete | 93.97 (100th) |
| GCA_002111555.1 | *Lonsdalea* | *populi* CFCC11748 | Contig | 91.09 (100th) |
| GCA_003269855.1 | *Lonsdalea* | *populi* CFCC11200 | Contig | 100 (100th) |
| GCA_003666245.1 | *Brenneria* | *alni* NCPPB 3934 | Contig | 100 (100th) |
| GCA_000406125.1 | *Brenneria* | *rubrifaciens* 6D370 | Complete | 100 (100th) |

### 8 Exploration of CAZy family frequencies

#### SI Table 20: Genus specific families in *Pectobacteriaceae*

CAZy families that are only found in one genus of *Pectobacteriaceae*, or only found in soft or hard plant tissue targeting genomes are listed in table 1.

Table 1: CAZy families found in only one genus, or only soft plant tissue targeting genomes, or hard plant tissue targeting genomes in *Pectobacteriacea*

| Genus | GH | GT | CE | PL | AA | CBM |
|---|---|---|---|---|---|---|
| *Pectobacterium* | GH121, GH146, GH18 | GT101, GT102, GT11, GT111, GT14, GT24, GT52 | | PL11 | AA10 | CBM13 |
| *Dickeya* | GH113, GH148, GH25, GH91 | GT97 | CE2 | PL10 | | CBM4 |
| **Soft tissue** | GH16, GH18, GH25, GH91, GH113, GH121, GH146, GH148 | GT11, GT14, GT24, GT25, GT52, GT97, GT101, GT102, GT111 | CE2 | PL10, PL11, PL35 | AA10 | CBM0, CBM4, CBM13, CBM91 |
| *Brenneria* | GH106 | GT21 | | PL17 | | |
| *Acerihabitans* | GH127, GH15 | | | | | |
| **Hard tissue** | GH15, GH37, GH39, GH51, GH106, GH127, GH140 | GT20, GT21, GT28 | | PL17 | | |

#### SI Figure 11: Core CAZome Families

The number of unique protein accessions was calculated for each CAZy family, per genome, and the mean number of proteins per CAZy family that was found in every genome (the "core CAZome") was plotted as a proportional area plot, breaking down the data per genus.

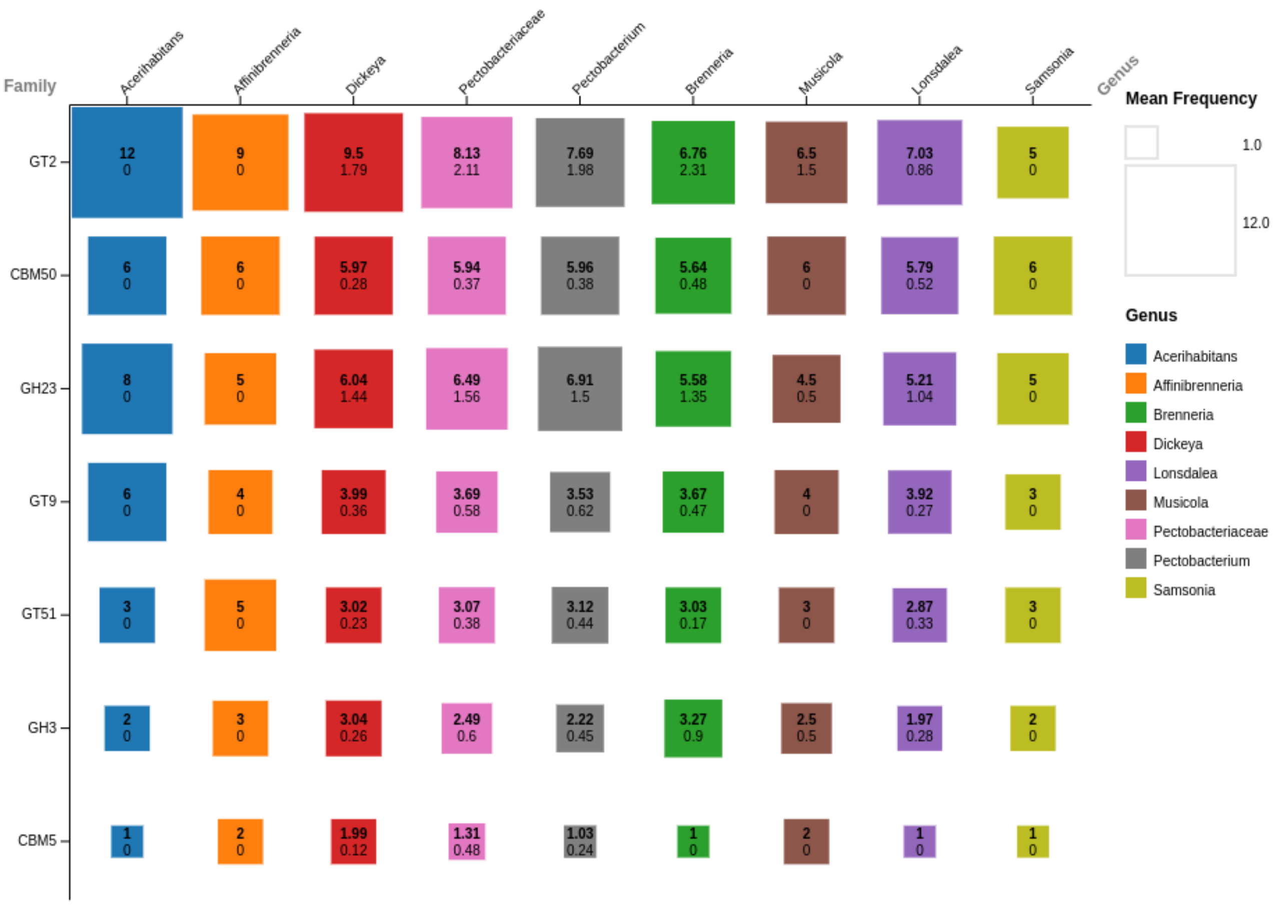


Figure 11: The core CAZome of *Pectobacteriaceae*. Proportional area plot of the mean number of CAZymes per CAZyme family that are present in all *Pectobacteriaceae* genomes (i.e. the core CAZome). Each square is annotated with the mean frequency (in bold), and standard deviation.

### 9 Coinfinder output

**SI Figure 12: Output from `coinfinder` with taxonomic annotations included (on the next page)**

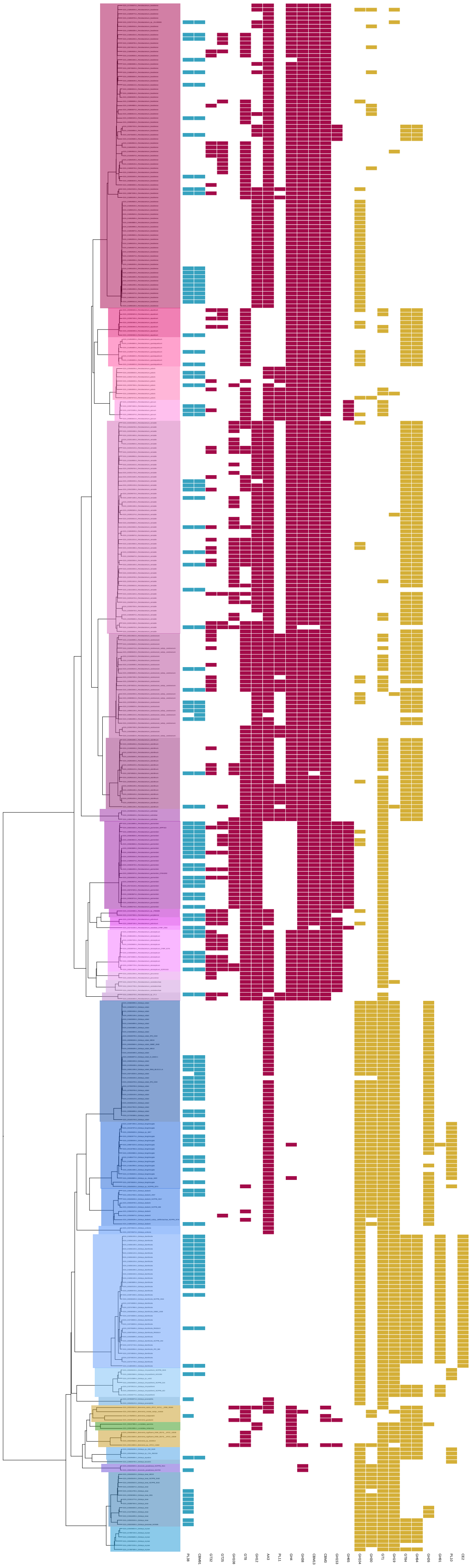
PL38
CBM91
GT32
GT25
GH108
GT8
GH12
AA3
PL11
GH4
GH38
CBM13
CBM3
GH153
GH65
GH104
GH30
GT1
GH19
GT84
GH94
GH26
GH91
PL10
CE2

#### SI Figure 13: ANI versus CAZyme family frequency tanglegram

The number of unique protein accessions was calculated for each CAZy family, per genome. These CAZy family frequencies were used to build a dendrogram. An ANI-based tree was built using all genomic sequences and `pyani`.

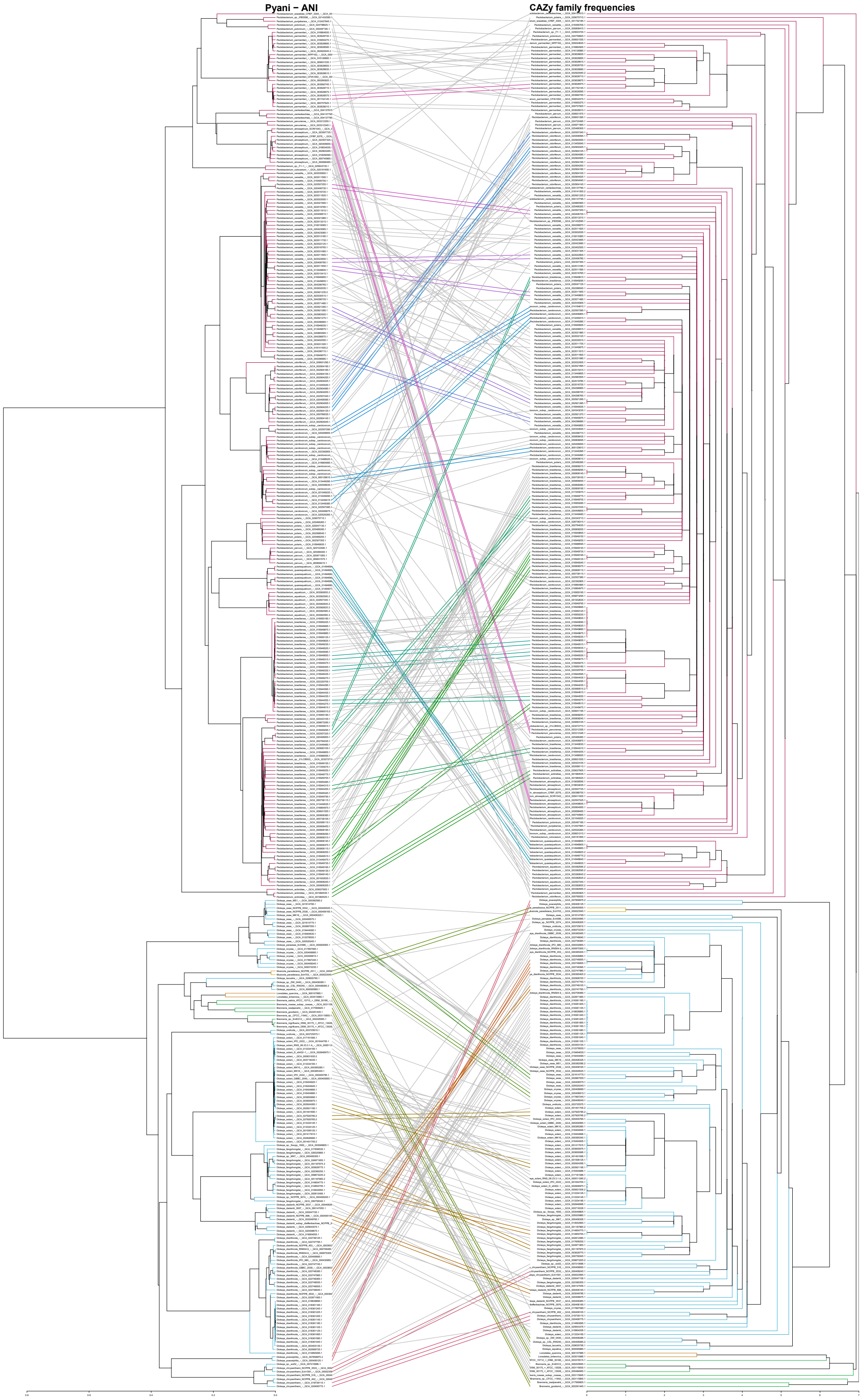


Figure 12: Tanglegram comparing the topologies of dendrograms generated from single linkage hierarchical clustering of CAZyme family frequencies and average nucleotide identities (ANI). *Leaf branches are colour coded by the genus classification of the corresponding genomes: pink, Pectobacterium; green, Brenneria; purple, Affinibrenneria; teal, Acerihabitans; gold, Musicola; blue, Dickeya; orange, Lonsdalea. Matching subtrees are indicated by colour-coded joining lines, matching the corresponding genomes between the dendrograms.*
